# Low-Coverage Genome Sequencing Outperforms Target Enrichment Phylogenomics

**DOI:** 10.64898/2026.06.05.730492

**Authors:** Michael G. Branstetter, Felipe V. Freitas, Ligia R. Benavides Silva, Silas Bossert, Bryan N. Danforth, Elizabeth A. Murray

## Abstract

Genome-scale data have transformed phylogenetic inference, yet most studies continue to rely on reduced-representation approaches that target a subset of loci to reduce cost and increase taxon sampling. Although effective, these methods require specialized laboratory workflows, constrain long-term data reuse, and may perform poorly with degraded DNA. Low-coverage whole genome sequencing (lcWGS) offers a streamlined alternative: shallow to moderate sequencing of complete genomes followed by bioinformatic extraction of loci of interest. Despite its promise, lcWGS has not been rigorously benchmarked against targeted enrichment using historical museum specimens. Here, we directly compared lcWGS and ultraconserved element (UCE) target enrichment across taxonomically diverse bee specimens collected between 1934 and 2021. Both data types were generated from the same Illumina libraries, enabling a controlled, head-to-head evaluation. Using standard UCE analytical pipelines, we quantified locus recovery, gene-tree support, and phylogenetic performance across sequencing methods and specimen age classes. We further assessed recovery of additional marker classes, including mitogenomes, BUSCO loci, and UCEs from a newly-designed, expanded probe set. Across all age categories, lcWGS consistently outperformed target enrichment, recovering more UCE loci and substantially longer alignments, with the largest gains observed in highly degraded specimens. Gene trees derived from lcWGS exhibited higher mean bootstrap support and greater topological concordance, translating into improved species-tree inference. In addition, lcWGS enabled recovery of markedly more non-target loci, expanding analytical flexibility beyond the original marker set. These results demonstrate that lcWGS not only matches but frequently exceeds the performance of targeted enrichment in museum-based phylogenomics, while providing broader genomic utility. As sequencing costs continue to decline, lcWGS represents a robust and forward-looking strategy for phylogenetic research, particularly in taxa with modest genome sizes and challenging DNA quality.

## Introduction

Advances in high-throughput sequencing, laboratory protocols, and computational power since the early 2010s have transformed molecular systematics and accelerated the transition to phylogenomics (Lemmon and Lemmon 2013, McCormack et al. 2013, Gillung et al. 2018). At the same time, short-read sequencing approaches that do not rely on locus-specific PCR have made it possible to recover highly fragmented DNA, expanding access to genomic data from historical museum specimens. This convergence of technological advances has fueled the rise of “museomics”, dramatically increasing the evolutionary and ecological value of natural history collections (Raxworthy and Smith 2021). Improved DNA extraction, library preparation, and sequencing protocols now routinely generate usable data from specimens collected more than a century ago, including small-bodied insects and rare or extinct taxa (Blaimer et al. 2016, Mikheyev et al. 2017, Schweizer et al. 2025).

Despite these advances, uncertainty remains regarding best practices for generating phylogenomic data from specimens of varying age and DNA quality. Most contemporary phylogenomic studies rely on reduced-representation approaches that sequence hundreds to thousands of loci rather than complete genomes. These methods—often termed target enrichment, transcriptomics, or RADseq-based approaches—reduce per-sample costs and computational burden while providing robust phylogenetic signal. Among arthropods and many other taxa, enrichment of ultraconserved elements (UCEs) has become especially widespread due to standardized laboratory workflows, scalable and publicly available bioinformatic pipelines, and demonstrated performance across deep and shallow evolutionary timescales (Baca et al. 2017, Branstetter et al. 2017a, Branstetter et al. 2017b, Van Dam et al. 2017, Branstetter and Longino 2019, Hedin et al. 2019, Almeida et al. 2023). Importantly, UCEs can be recovered from highly degraded DNA, with successful applications to century-old arthropod specimens (Blaimer et al. 2016, Wood et al. 2018, Freitas et al. 2023). More recently, Goodman et al. (2023) demonstrated that targeted enrichment performs effectively across odonate museum specimens spanning a broad temporal range, further highlighting the growing utility of enrichment-based phylogenomics for historical insect collections.

As sequencing costs continue to decline, whole-genome sequencing (WGS) is increasingly considered as an alternative to reduced-representation methods. Although historically cost-prohibitive, WGS offers several advantages: simplified laboratory workflows that do not require lineage-specific probe design, independence from prior genomic resources, and the generation of broadly reusable datasets suitable for diverse downstream analyses (Allen et al. 2017). To mitigate costs, many studies sequence genomes at moderate depth and extract loci of interest bioinformatically—a strategy often referred to as genome skimming or low-coverage whole-genome sequencing (lcWGS) (Zeng et al. 2018, Pezzini et al. 2023, Taite et al. 2023, Hu et al. 2024, Chen et al. 2025). While genome skimming has traditionally been used to recover organellar genomes and high-copy nuclear markers, recent work demonstrates that lcWGS can also recover non-repetitive nuclear loci, including UCEs and single-copy orthologs such as BUSCO genes (Zhang, Feng et al. 2019a).

Despite its promise, the relative performance of lcWGS and target enrichment remains poorly evaluated, particularly for historical museum specimens. UCE enrichment has proven effective with degraded insect DNA and across museum age gradients (Goodman et al. 2023), yet it remains unclear whether hybrid capture or lcWGS is more resilient to DNA fragmentation associated with specimen age. Critically, direct, controlled comparisons of these approaches using the same museum specimens and sequencing libraries are lacking.

Here, we conduct a head-to-head comparison of lcWGS and UCE target enrichment using taxonomically diverse bee specimens collected across almost 90 years and representing all major bee families. From identical Illumina libraries, we either sequenced whole genomes at moderate depth (∼20× coverage) or enriched libraries for UCE loci prior to sequencing, enabling a controlled methodological benchmark (Fig. 1). Although “low coverage” is variably defined in the literature, we use the term here relative to typical high-depth genome projects (≥100× coverage), with the goal of recovering phylogenetic markers rather than assembling chromosome-level genomes. We do not attempt to determine the minimum coverage required for comparable results, leaving that optimization question for future study.

**Figure 1.**
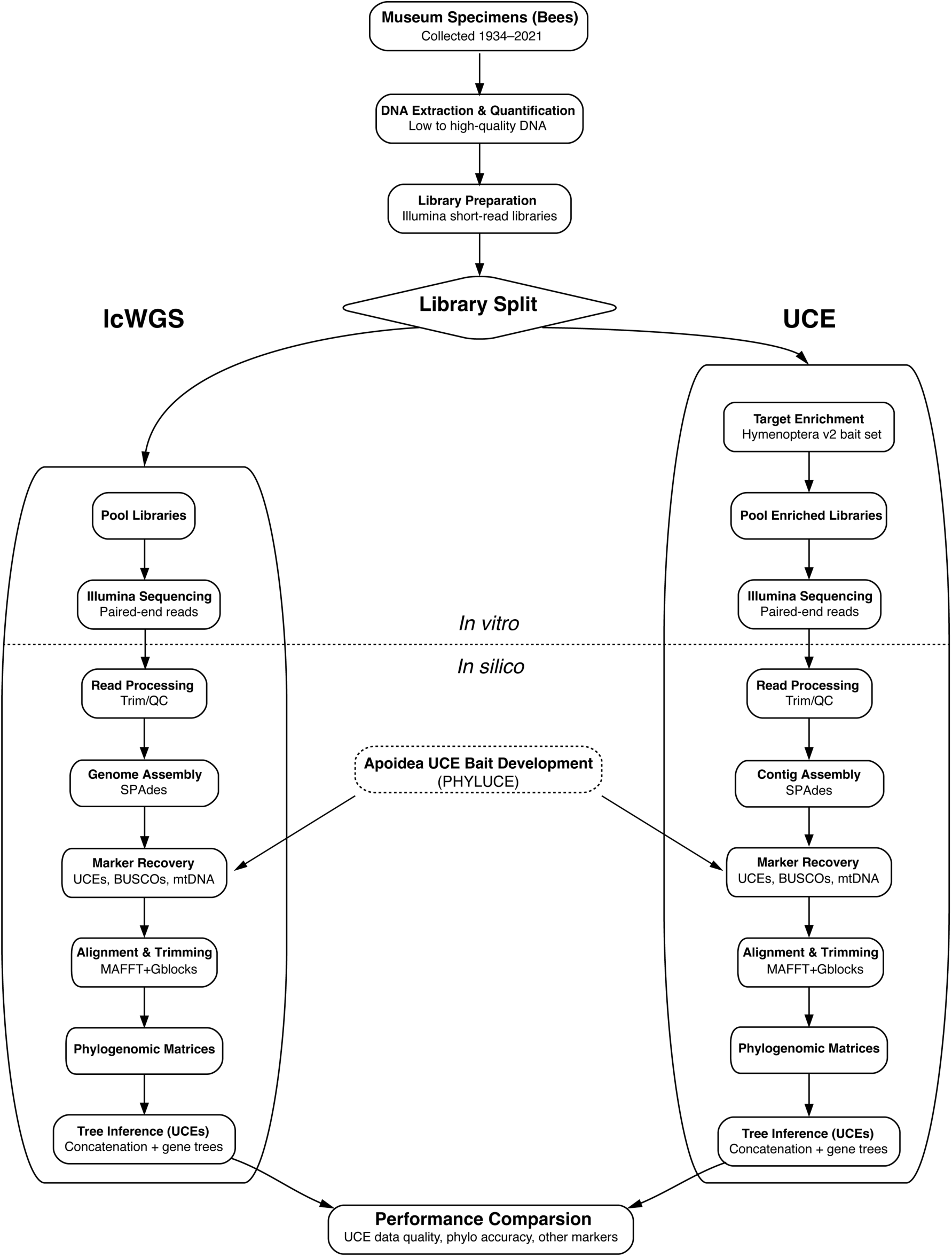
Project flow chart comparing lab and data processing workflows for the low-coverage, whole genome sequencing (lcWGS) and ultraconserved element (UCE) datasets. Both workflows used the same Illumina libraries.

We processed both datasets using comparable bioinformatic workflows and evaluated performance across multiple dimensions, including recovery of UCE loci (number and length), recovery of additional marker classes (BUSCO genes, mitochondrial genomes, and a novel expanded Apoidea-specific UCE probe set), and phylogenetic resolution under both concatenation and gene-tree species-tree frameworks. By benchmarking marker recovery and phylogenetic performance across specimen age classes, we provide the first systematic assessment of lcWGS versus target enrichment in museum insects.

## Materials & Methods

### Specimen Sampling

We sampled 60 bee museum specimens representing all major bee families and spanning collection years from 1934 to 2021. Specimens were selected to include multiple representatives from each family and from four age categories: (1) pre-1960, (2) 1960–1979, (3) 1980–1999, and (4) 2000–2021. The final dataset included 10 Andrenidae, 15 Apidae, 10 Colletidae+Stenotritidae, 9 Halictidae, 8 Megachilidae, and 8 Melittidae. Age categories contained 6, 15, 15, and 24 specimens, respectively (Table 1).

**Table 1.**
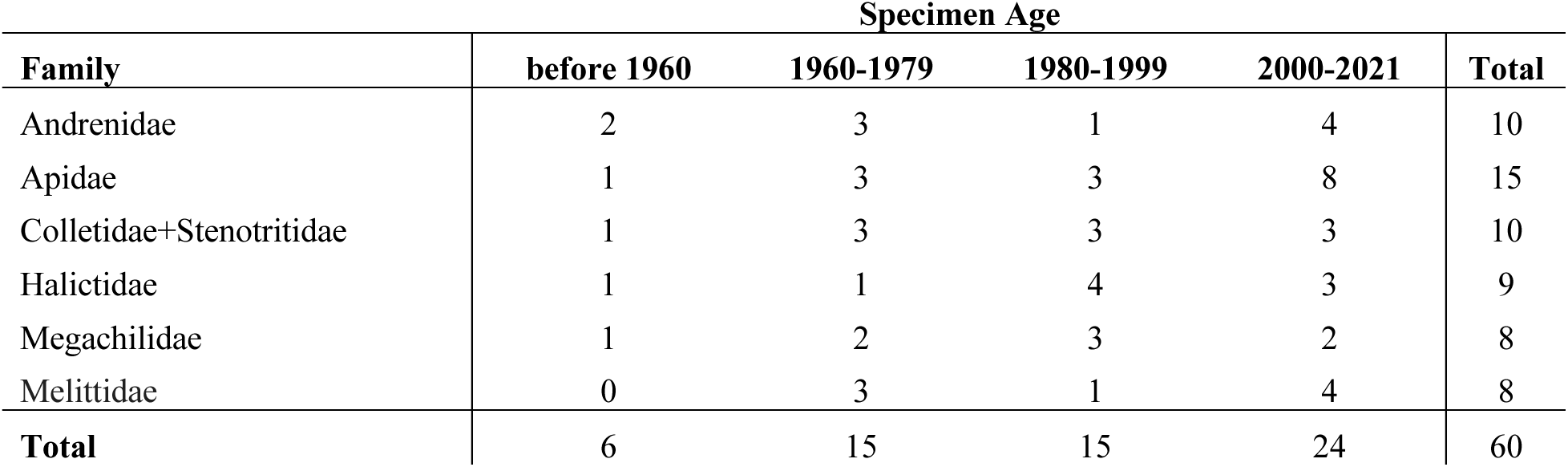
Counts of specimens by bee family and age category. The families Colletidae and Stenotritidae were combined into a single group.

| Family | Specimen Age |  |  |  | Total |
| --- | --- | --- | --- | --- | --- |
|  | before 1960 | 1960-1979 | 1980-1999 | 2000-2021 |  |
| Andrenidae | 2 | 3 | 1 | 4 | 10 |
| Apidae | 1 | 3 | 3 | 8 | 15 |
| Colletidae+Stenotritidae | 1 | 3 | 3 | 3 | 10 |
| Halictidae | 1 | 1 | 4 | 3 | 9 |
| Megachilidae | 1 | 2 | 3 | 2 | 8 |
| Melittidae | 0 | 3 | 1 | 4 | 8 |
| **Total** | 6 | 15 | 15 | 24 | 60 |

Sample size was determined based on the output capacity of a single Illumina NovaSeq 6000 S4 lane and a target sequencing depth of ∼20× per specimen for whole-genome sequencing. Because genome assemblies are unavailable for most focal species, coverage was estimated assuming a mean bee genome size of ∼500 Mb (based on data in NCBI accessed in November 2021). Complete specimen metadata and voucher information are provided in Table S1.

### Library Preparation

Genomic DNA was extracted using Zymo Quick-DNA MiniPrep Plus kits (Zymo Research, Irvine, CA, USA) with the following modifications: (a) overnight Proteinase-K digestion, (b) pre-heating the elution buffer to 55°C, and (c) two elutions of 60 μL each. For most specimens, one to three legs were removed using sterile technique. For several smaller taxa, whole-body non-destructive extractions were performed; specimens were subsequently rinsed in 95% ethanol and remounted.

DNA concentration was quantified with a Qubit 3.0 fluorometer (Thermo Fisher Scientific), and fragment size distributions were assessed with a TapeStation 4150 (Agilent Technologies). Up to 70 ng of DNA was used for Illumina library preparation with KAPA Hyper Prep kits (Roche Sequencing Solutions) and custom 8 bp dual-index adapters (Glenn et al. 2019). DNA was sheared for 0–120 seconds using a QSonica Q800 R3 sonicator, depending on initial fragment size (200 bp–5.6 kb). For higher-quality samples, we targeted a mean fragment size of ∼400 bp. Libraries were cleaned with 3× SPRI beads (Rohland and Reich 2012) and amplified with 12 PCR cycles. Final libraries were assessed with Qubit and TapeStation.

### Sequencing Design: lcWGS and UCE Enrichment

Libraries were pooled into 6 groups of 10 samples based on DNA concentration measured by Qubit. Aliquots from each pool were processed using two workflows: (1) low-coverage whole-genome sequencing (lcWGS) and (2) ultraconserved element (UCE) target enrichment (see Fig. 1 flow chart outlining project design).

#### lcWGS

Pools were quantified using qPCR (Applied Biosystems ViiA 7) and combined into a single sequencing pool. Adapter dimers were removed by BluePippin size selection (250–900 bp), followed by 3× SPRI cleanup. The final pool was sequenced on one NovaSeq 6000 S4 lane by Novogene in Sacramento, CA (Novogene Corporation Inc.). We aimed for ∼20× sequencing coverage per sample and consider this to be “low coverage” relative to conventional genome projects targeting ≥100× Illumina depth for high-contiguity assemblies.

#### UCE Target Enrichment

Each of the six initial pools was enriched using the ant-bee-specific bait set (Hymenoptera 2.5Kv2AB) (Grab et al. 2019), a subset of the principal bait set (Hymenoptera 2.5Kv2P) (Branstetter et al. 2017b). This kit includes 9,068 baits targeting 2,545 UCE loci plus legacy exonic markers and is commercially available from Daicel Arbor Biosciences (Ann Arbor, MI, U.S.A.).

Enrichment followed Arbor Biosciences v3.02 protocol for day 1 and a standard UCE protocol for day 2 (UCE Enrichment v1.5 available at http://s3.ultraconserved.org/protocols/illumina-seqcap-hybridization-with-myselect.pdf) (Faircloth et al. 2012). Hybridization was conducted at 65°C for 24 h. On-bead PCR amplification used 18 cycles. Enriched pools were cleaned (1× SPRI), quantified by qPCR, and combined into two final pools (30 samples each). Sequencing was performed by Novogene on a NovaSeq 6000 partial lane, targeting ∼1 Gb per sample (∼61 Gb total), a depth typically sufficient for high-quality UCE recovery and consistent with recent studies. Laboratory metadata are available in Table S2. All raw sequence data have been deposited in the NCBI Sequence Read Archive (accession numbers in Table S3).

### Read Processing and Assembly

Raw lcWGS reads were demultiplexed using the *demuxbyname2* program, which is part of the BBTools v39.00 package (Bushnell 2014). Read counts and quality were assessed with FastQC v0.12.1 (Andrews 2010) and summarized with MultiQC v1.26 (Ewels et al. 2016). To ensure comparability, lcWGS and UCE datasets were processed using identical trimming and assembly workflows where applicable, with most steps run within the PHYLUCE v1.7.1 software environment (Faircloth 2016). Reads were trimmed using Illumiprocessor v2.10 (Faircloth 2013) with sample-specific adapter sequences. Genome size estimates were generated using GenomeScope2 (Ranallo-Benavidez et al. 2020). Assemblies were performed with SPAdes v3.14.1 (Bankevich et al. 2012) via PHYLUCE wrappers and default parameter settings. Sequencing coverage of the initial, bulk assemblies and contigs containing UCE loci, was assessed using the PHYLUCE mapping workflow which aligns trimmed reads to contigs and computes per-base pair coverage.

Five samples with exceptionally high read counts failed assembly due to computational memory limitations rather than insufficient data. For these samples, reads were normalized to 40× coverage using *bbnorm* (BBTools) prior to assembly, retaining depth similar to enrichment datasets.

UCE-enriched reads were processed from read trimming to assembly identically to lcWGS reads; except we did not estimate genome size using GenomeScope2 given that the data should not be representative of the entire genome.

### UCE Locus Extraction and Analysis

UCE loci were extracted from assemblies using the PHYLUCE “Harvesting UCE loci from genomes” tutorial workflow (https://phyluce.readthedocs.io/en/latest/tutorials/tutorial-3.html). Contigs were converted to 2bit format using faToTwoBit from the UCSC Genome Browser Group’s suite of programs (Kent 2002), and bait sequences from the Hymenoptera v2.5Kv2P bait set were mapped with minimum thresholds of 80% coverage and 80% identity. Loci were extracted with 1 kb flanking regions. Loci were matched to probes (*phyluce_assembly_match_contigs_to_probes*) using the Hymenoptera v2.5Kv2P bait set, aligned with MAFFT v7.475 (Katoh and Standley 2013), trimmed using Gblocks (Talavera and Castresana 2007) (reduced stringency: b1=0.5, b2=0.5, b3=12, b4=7), and filtered for 75% taxon occupancy. Concatenated supermatrices were then constructed for phylogenetic analyses. To ensure comparability, assembled contigs from the UCE enriched dataset were processed starting from *phyluce_assembly_match_contigs_to_probes* using identical parameters.

Per-sample UCE performance metrics were calculated and included: (1) number of loci recovered, (2) mean locus length, and (3) number of loci >1 kb. These metrics were calculated by exploding the monolithic UCE FASTA file generated by PHYLUCE by taxon and then running the PHYLUCE script *phyluce_assembly_get_fasta_lengths*. The UCE loci were assessed prior to alignment or trimming for downstream phylogenetic analyses.

### Phylogenetic Inference

To assess phylogenetic accuracy of the UCE data, we performed concatenated and gene tree-based analyses. Concatenated datasets were analyzed with IQ-TREE v2.3.5 (Minh, B. Q. et al. 2020b) under GTR+F+G4 without partitioning and support was assessed with 1,000 ultrafast bootstrap replicates (UFB) (Minh, B. Q. et al. 2013, Hoang et al. 2018). Gene trees were inferred per locus using IQ-TREE with full model selection (-m MFP -AICc) and 1,000 UFB replicates.

To assess the quality of concatenated trees, we compared topologies and support values while also considering established understanding of bee phylogenetic relationships (Almeida et al. 2023). To assess the quality of gene trees, we examined mean UFB support and phylogenetic accuracy. We operationally defined gene-tree accuracy as quartet similarity to a reference topology (Asher et al. 2022), calculated using the R package Quartet (Smith 2019). This metric quantifies agreement among quartets defined over the subset of taxa shared between each gene tree and the reference topology, thereby accommodating missing taxa across loci. Gene concordance factors (gCF) (Minh, Bui Quang et al. 2020a) were also calculated in IQ-TREE v3 (Wong et al. 2025). Both lcWGS and UCE concatenated tree topologies were used as reference topologies for the gene-tree accuracy analyses to avoid reference bias and account for any topological differences.

### Additional Marker Recovery

To further compare lcWGS data to UCE-enriched data, we extracted three additional types of markers from the lcWGS and UCE datasets: Benchmarking Universal Single Copy Orthologs (or BUSCOs), mitogenomes, and a novel, expanded UCE locus set designed from a more inclusive set of taxa than the full Hymenoptera v2.5Kv2P bait set.

#### BUSCO

To evaluate assembly quality and quantify the number of recovered BUSCO loci found in each dataset, we used BUSCO v5 (Manni et al. 2021) and BlobTools v2 (Laetsch and Blaxter 2017). For the focal BUSCO database, we used hymenoptera_odb10, which includes 5,991 single copy orthologs; however, we also examined results for eukaryota_odb10, arthropoda_odb10, insecta_odb10, and endopterygota_odb10 gene sets, which include 255, 1013, 1367, and 2124 single copy genes, respectively. BlobTools was used to generate genome assembly snail plots and to assess the data for non-target sequence contamination.

#### Mitogenomes

Mitogenomes were extracted using MitoFinder v1.4.1 (Allio et al. 2020) with 244 hymenopteran mitogenomes downloaded from NCBI as references (accessed 13 June 2025). MitoFinder was run with the following parameter settings: *-o 5 -p 8 -m 24 -e 0.001 -n 15 - -min-contig-size 500 --blast-size 20 --ignore*. For each sample, we examined two metrics for comparison: (1) the total number of mtDNA genes recovered, and (2) the length of the largest mtDNA contig.

#### Expanded UCE Set

We explored if it was possible to extract a larger set of UCE loci from the assemblies by first identifying an expanded set of UCE loci present in Apoidea (bees and apoid wasp relatives), a subclade within Hymenoptera. By searching for conserved regions within a shallower clade of taxa we reasoned that it should be possible to identify a larger number of UCE loci given that taxa are more closely related and share a more recent common ancestor. To accomplish this, we selected a set of 13 Apoidea reference genomes available from NCBI including 11 bees (Table S4). We then followed PHYLUCE tutorial four (https://phyluce.readthedocs.io/en/latest/tutorials/tutorial-4.html), which outlines how to identify UCE loci across genomes and design baits to enrich these loci, both for *in vitro* and *in silico* analyses. We selected *Macropis europaea* as the base genome for the analysis because it is a high-quality, chromosome-level assembly and it belongs to our target group. Reads were simulated from the remaining genomes using ART v2.5.8 (Huang et al. 2012) and aligned to the base genome using PHYLUCE scripts and the program LASTZ v1.04 (Harris 2007), resulting in the identification of genomic regions that have < 5% sequence divergence. We merged closely approximated regions of conservation and removed repetitive intervals. We then searched for locus presence across the genomes and retained the set of loci shared between the base genome and all 12 of the remaining genomes. This resulted in the identification of 20,832 conserved regions. For each conserved region, we extracted 160 bp sequences and designed baits at 3x tiling density to enrich each locus. We removed duplicates from the baits and aligned the resulting dupe-free baits to the exemplar genomes. We buffered the regions to 180 bp and sliced out resulting sequences into FASTA format. We examined the number of loci shared amongst the exemplar taxa and we settled upon loci present in all 12 taxa. A final bait set was designed from these loci at 3x tiling density and duplicates were removed from the final bait set.

We refer to this final bait set as “Apoidea 20Kv1P”. After bait design, we followed the same align-and-slice workflow described above and extracted Apoidea 20Kv1P UCE loci from each of the two datasets. For this set of loci, we report on the number of extracted UCE loci as a primary metric, along with stats for mean UCE length and number of contigs > 1kb, like the Hymenoptera 2.5Kv2P UCE datasets. To evaluate phylogenetic accuracy of the new set of loci, we estimated a concatenated phylogenetic tree using IQ-TREE v2 and the UCEs recovered from the lcWGS dataset. Loci were processed for phylogenetic analysis following the same workflow described above.

### Statistics

Most comparative statistical analyses were performed in R v4.5.1 (Team 2019), with plots generated with ggplot2 (Wickham 2016). Relationships among sequencing metrics and collection year were examined with ordinary least squares linear regression using *lm()*. We extracted slope, intercept, 𝑅^!^, and 𝑝-values and plotted fitted lines. Collection year was treated as a proxy for specimen/DNA quality.

We examined read counts by age category for each dataset (lcWGS and UCE) using violin plots and tested for significant differences among age categories using a Kruskal-Wallis test. We then performed pairwise Wilcoxon tests to compare between age categories. We adjusted p-values for multiple testing using Benjamini–Hochberg (FDR) and additionally report Bonferroni-adjusted values in the results table.

To compare lcWGS and UCE datasets for each sample-based metric (e.g. # of UCEs, mean UCE length, # of contigs > 1 kb), we generated violin plots and/or boxplots and performed paired tests. We tested normality of the paired differences (lcWGS − UCE) with the Shapiro–Wilk test; when normality held (p > 0.05) we used a two-sided paired t-test, otherwise a two-sided Wilcoxon signed-rank test. We repeated these paired comparisons for all samples combined and for each age category separately. We adjusted p-values for multiple testing using Benjamini–Hochberg (FDR) (stars reflect BH-adjusted p-values) and additionally report Bonferroni-adjusted values in the results table. For the gene tree metrics (e.g. mean bootstrap, phylogenetic accuracy, gCF), we visualized the values for each dataset as violin plots and checked for statistical differences using the Wilcoxon test and p-values adjusted for multiple comparisons. Statistical significance on plots is annotated as * 𝑝 < 0.05, ** 𝑝 < 0.01, and *** 𝑝 < 0.001 (BH-adjusted where applicable).

## Results

### Sequencing

For the lcWGS dataset, we recovered a mean of 45.7 million raw paired-end (PE) reads per sample (range: 17.6–87.5 M PE reads; Table S5). After trimming, a mean of 44.3 M PE reads remained (range: 17.2–79.1 M). In the UCE-enriched dataset, we recovered a mean of 4.1 M raw PE reads per sample (range: 0.7–13.0 M) and 4.0 M trimmed PE reads (range: 0.7–12.2 M).

There was a significant negative correlation between collection year and raw PE read count in the lcWGS dataset (Fig. S1; y = −199,777.87x + 443,222,135.27; r² = 0.092; p = 0.0185). In contrast, the UCE dataset showed a significant positive correlation (y = 46,326.18x − 88,056,657.26; r² = 0.158; p = 0.00164). However, slopes were shallow in both cases, and collection year explained only a small proportion of the total variance. When samples were grouped by age category, no significant differences in read counts were detected within either dataset (Fig. S1, Table S6; pairwise Wilcoxon tests with Benjamini–Hochberg correction; p > 0.05). Results were similar for the trimmed reads (Fig. S2).

GenomeScope2 estimated a mean genome size of 332.5 Mb (95% CI 280.7-384.3 Mb), with the range extending from 10,152 bp (*Centris obscurior*) to 769.8 Mb (*Centris rhodopus*) (Fig. S3). Ten samples had estimated genome sizes below 100 Mb (*Calliopsis zebrata, Centris obscurior, Eucera nigrescens, Eufriesea engeli, Nogueirapis butteli, Notoxaea ferruginea, Panurginus ineptus, Protandrena angusticeps, Protoxaea gloriosa, Stenotritus greavesi*), likely indicating erroneous genome size estimates; most remaining samples, however, returned values consistent with previously reported bee genome size estimates. We detected no significant relationship between raw read counts and genome size estimates (Fig. S3; y=1.099x+282.247; r^2^ = 0.006; p = 0.542).

Sequencing coverage was estimated for both lcWGS and UCE-enriched datasets using complete assembled contigs (bulk contigs) and for contigs containing UCE loci (Fig. S4). For the lcWGS dataset, mean coverage was 23.3× for bulk contigs and 42.1× for UCE-containing contigs. In the UCE-enriched dataset mean coverage was 9.46 and 67.08. The magnitude of the difference between bulk and UCE contig coverage for the two datasets was similar across age categories, and in all comparisons UCE-containing contigs exhibited significantly higher coverage than bulk contigs.

### UCE Recovery and Length

To compare dataset performance, we examined (1) the number of extracted UCE loci, (2) mean UCE locus length, and (3) the number of UCE contigs >1 kb. Across all samples, the lcWGS dataset outperformed the UCE-enriched dataset for all three metrics (Fig. 2; Table S5). For the lcWGS dataset, we recovered significantly more loci (mean: 2,355 vs. 2,170; p < 0.001; Wilcoxon signed-rank test), significantly longer loci (1,704 bp vs. 988 bp; p < 0.001), and substantially more UCE contigs >1 kb (2,038 vs. 877; p < 0.001). When stratified by age category, this pattern was largely consistent: lcWGS exceeded UCE enrichment in nearly all comparisons, with all but two tests being statistically significant (Fig. 2; Table S5).

**Figure 2.**
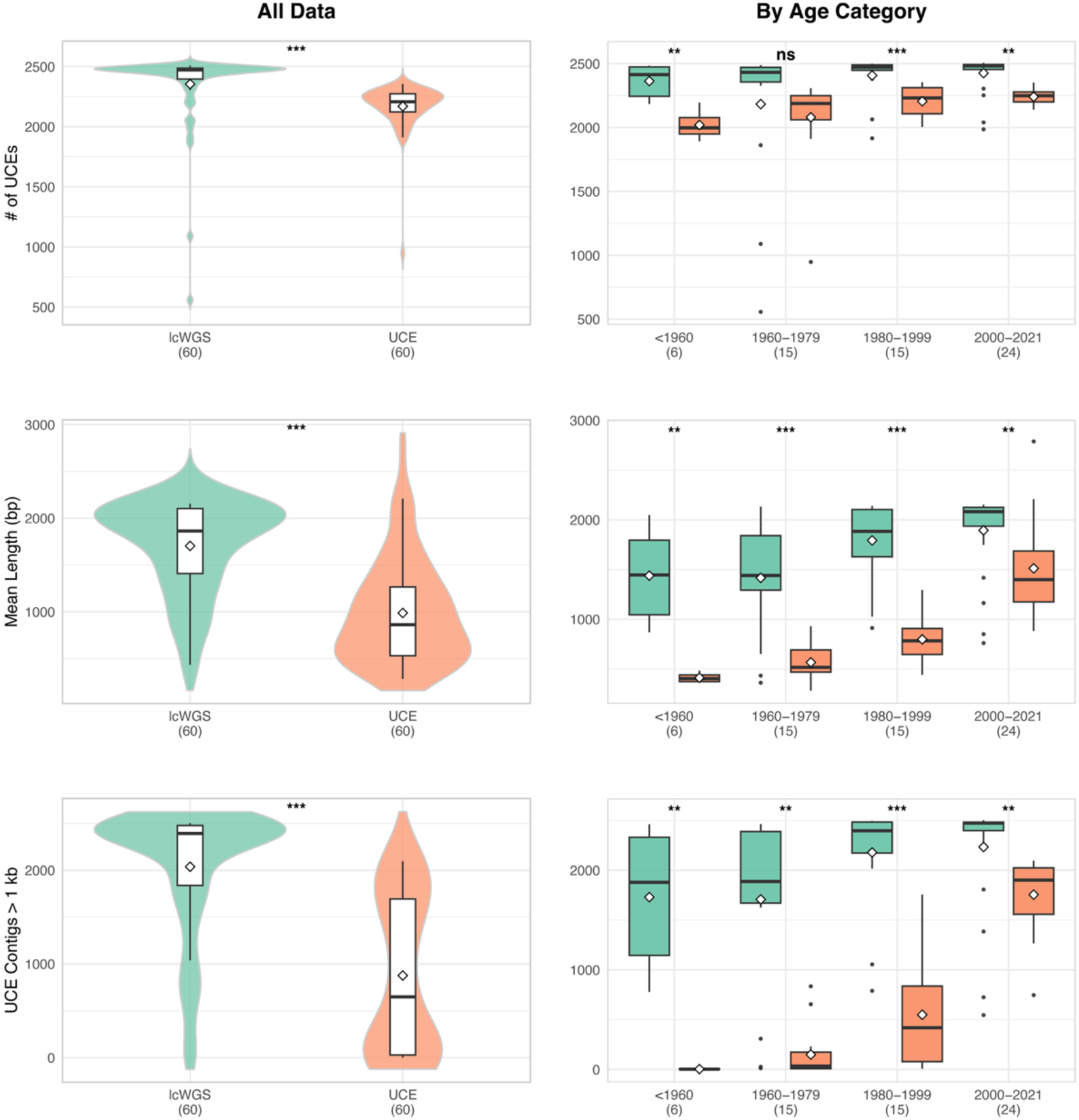
Comparison of UCE recovery metrics between lcWGS and UCE-enriched datasets. Left column shows all samples combined; right column shows results separated by age category. From top to bottom: (1) number of extracted UCE loci, (2) mean UCE locus length (bp), and (3) number of UCE contigs >1 kb. Violin plots show distribution density, boxplots show medians and interquartile ranges. Asterisks denote significance of paired comparisons between lcWGS and UCE datasets (Wilcoxon signed-rank or paired t-test as appropriate; Benjamini–Hochberg adjusted p-values): * p < 0.05, ** p < 0.01, *** p < 0.001. Sample sizes (n) are indicated on the x-axis. All comparisons are paired by sample.

Both datasets showed trends between collection year and UCE recovery metrics (Fig. 3; Table 2). In the UCE-enriched dataset, all three metrics exhibited significant positive correlations with collection year, and r² values were consistently higher than those observed for lcWGS. This pattern indicates a stronger dependence of UCE-enriched performance on specimen age, particularly for mean locus length and the number of UCE contigs >1 kb. In contrast, for lcWGS data, only mean UCE length showed a significant relationship with collection year (p = 0.0132), and effect sizes were smaller overall.

**Figure 3.**
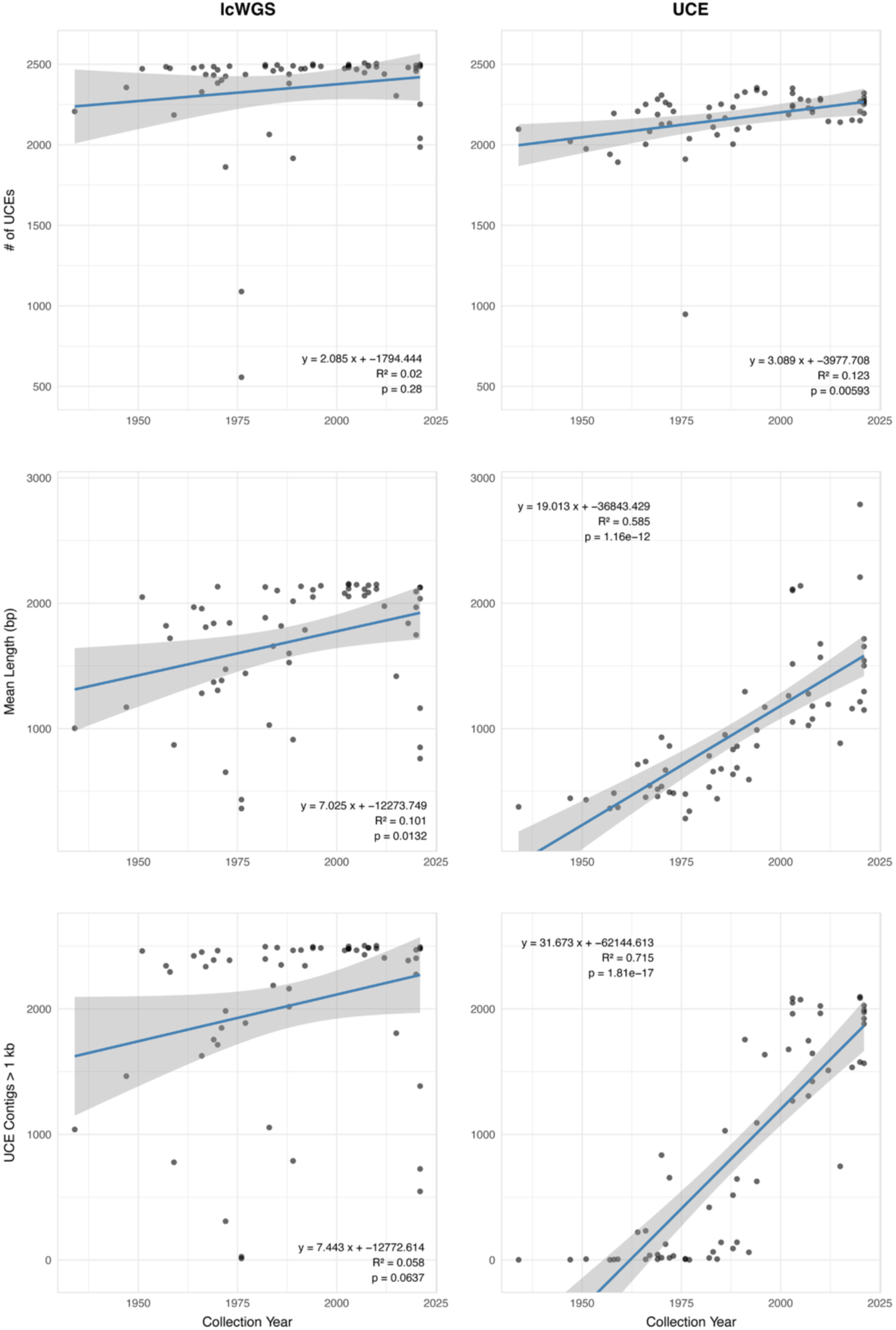
Relationships between UCE recovery metrics and specimen collection year. Left column shows lcWGS results; right column shows UCE-enriched results. From top to bottom: (1) number of extracted UCE loci, (2) mean UCE locus length (bp), and (3) number of UCE contigs >1 kb. Solid lines represent ordinary least squares regression fits. Regression statistics (slope, r², and p-values) are provided in each graph and Table 2.

**Table 2.** Statistical results comparing data quality metrics between the lcWGS and UCE-enriched datasets.

| Metric | Group | n | lcWGS mean | UCE mean | Mean difference | Test | Statistic | BH-adjusted p | Significance |
| --- | --- | --- | --- | --- | --- | --- | --- | --- | --- |
| # of UCEs | All | 60 | 2354.47 | 2169.57 | 184.9 | Wilcoxon signed-rank | 1578 | <0.001 | *** |
| # of UCEs | <1960 | 6 | 2363.17 | 2020.17 | 343 | Paired t-test | 5.324 | 0.004 | ** |
| # of UCEs | 1960-1979 | 15 | 2183.2 | 2079.93 | 103.27 | Wilcoxon signed-rank | 84 | 0.182 | ns |
| # of UCEs | 1980-1999 | 15 | 2407.8 | 2205.13 | 202.67 | Paired t-test | 5.029 | <0.001 | *** |
| # of UCEs | 2000-2021 | 24 | 2426 | 2240.71 | 185.29 | Wilcoxon signed-rank | 259.5 | 0.002 | ** |
| Mean Length (bp) | All | 60 | 1704.18 | 987.52 | 716.66 | Wilcoxon signed-rank | 1757 | <0.001 | *** |
| Mean Length (bp) | <1960 | 6 | 1438.9 | 412.3 | 1026.6 | Paired t-test | 5.346 | 0.004 | ** |
| Mean Length (bp) | 1960-1979 | 15 | 1416.95 | 567.29 | 849.66 | Paired t-test | 6.94 | <0.001 | *** |
| Mean Length (bp) | 1980-1999 | 15 | 1792.87 | 798.28 | 994.59 | Paired t-test | 11.097 | <0.001 | *** |
| Mean Length (bp) | 2000-2021 | 24 | 1894.59 | 1512.25 | 382.34 | Paired t-test | 3.598 | 0.002 | ** |
| UCE Contigs > 1 kb | All | 60 | 2037.87 | 877 | 1160.87 | Wilcoxon signed-rank | 1757 | <0.001 | *** |
| UCE Contigs > 1 kb | <1960 | 6 | 1729.83 | 3.33 | 1726.5 | Paired t-test | 5.774 | 0.003 | ** |
| UCE Contigs > 1 kb | 1960-1979 | 15 | 1707.27 | 149.47 | 1557.8 | Wilcoxon signed-rank | 120 | 0.001 | ** |
| UCE Contigs > 1 kb | 1980-1999 | 15 | 2178.53 | 549.13 | 1629.4 | Paired t-test | 10.404 | <0.001 | *** |
| UCE Contigs > 1 kb | 2000-2021 | 24 | 2233.58 | 1755.04 | 478.54 | Wilcoxon signed-rank | 247 | 0.006 | ** |
| BUSCO Hymenoptera | All | 60 | 3537.65 | 154.72 | 3382.93 | Wilcoxon signed-rank | 1830 | <0.001 | *** |
| BUSCO Hymenoptera | <1960 | 6 | 2401.83 | 4.5 | 2397.33 | Paired t-test | 3.346 | 0.022 | * |
| BUSCO Hymenoptera | 1960-1979 | 15 | 2369.27 | 30.93 | 2338.33 | Paired t-test | 5.56 | <0.001 | *** |
| BUSCO Hymenoptera | 1980-1999 | 15 | 3755.53 | 65.8 | 3689.73 | Wilcoxon signed-rank | 120 | 0.001 | ** |
| BUSCO Hymenoptera | 2000-2021 | 24 | 4415.67 | 325.21 | 4090.46 | Wilcoxon signed-rank | 300 | <0.001 | *** |
| # of mtDNA Genes | All | 60 | 13.08 | 8.05 | 5.03 | Wilcoxon signed-rank | 1091.5 | <0.001 | *** |
| # of mtDNA Genes | <1960 | 6 | 13.5 | 2.5 | 11 | Paired t-test | 10.333 | <0.001 | *** |
| # of mtDNA Genes | 1960-1979 | 15 | 13.13 | 5.6 | 7.53 | Paired t-test | 5.573 | <0.001 | *** |
| # of mtDNA Genes | 1980-1999 | 15 | 13.07 | 6.87 | 6.2 | Paired t-test | 6.254 | <0.001 | *** |
| # of mtDNA Genes | 2000-2021 | 24 | 12.96 | 11.71 | 1.25 | Wilcoxon signed-rank | 73 | 0.059 | ns |
| Apoidea UCEs | All | 60 | 12473.13 | 5014.25 | 7458.88 | Wilcoxon signed-rank | 1812 | <0.001 | *** |
| Apoidea UCEs | <1960 | 6 | 13221 | 1386 | 11835 | Paired t-test | 23.258 | <0.001 | *** |
| Apoidea UCEs | 1960-1979 | 15 | 12163.73 | 3018.47 | 9145.27 | Paired t-test | 11.489 | <0.001 | *** |
| Apoidea UCEs | 1980-1999 | 15 | 12774.53 | 4162.67 | 8611.87 | Paired t-test | 11.484 | <0.001 | *** |
| Apoidea UCEs | 2000-2021 | 24 | 12291.17 | 7700.92 | 4590.25 | Paired t-test | 5.778 | <0.001 | *** |

On an individual sample basis, lcWGS recovered fewer UCE loci than UCE enrichment in 8 of 60 specimens (Fig. 4). However, in five of these eight cases, lcWGS contigs had greater mean lengths (Table S4), partially offsetting the reduced locus count. Together, these results demonstrate that lcWGS consistently recovers more complete and longer UCE loci across a broad range of specimen ages, with reduced sensitivity to DNA degradation.

**Figure 4.**
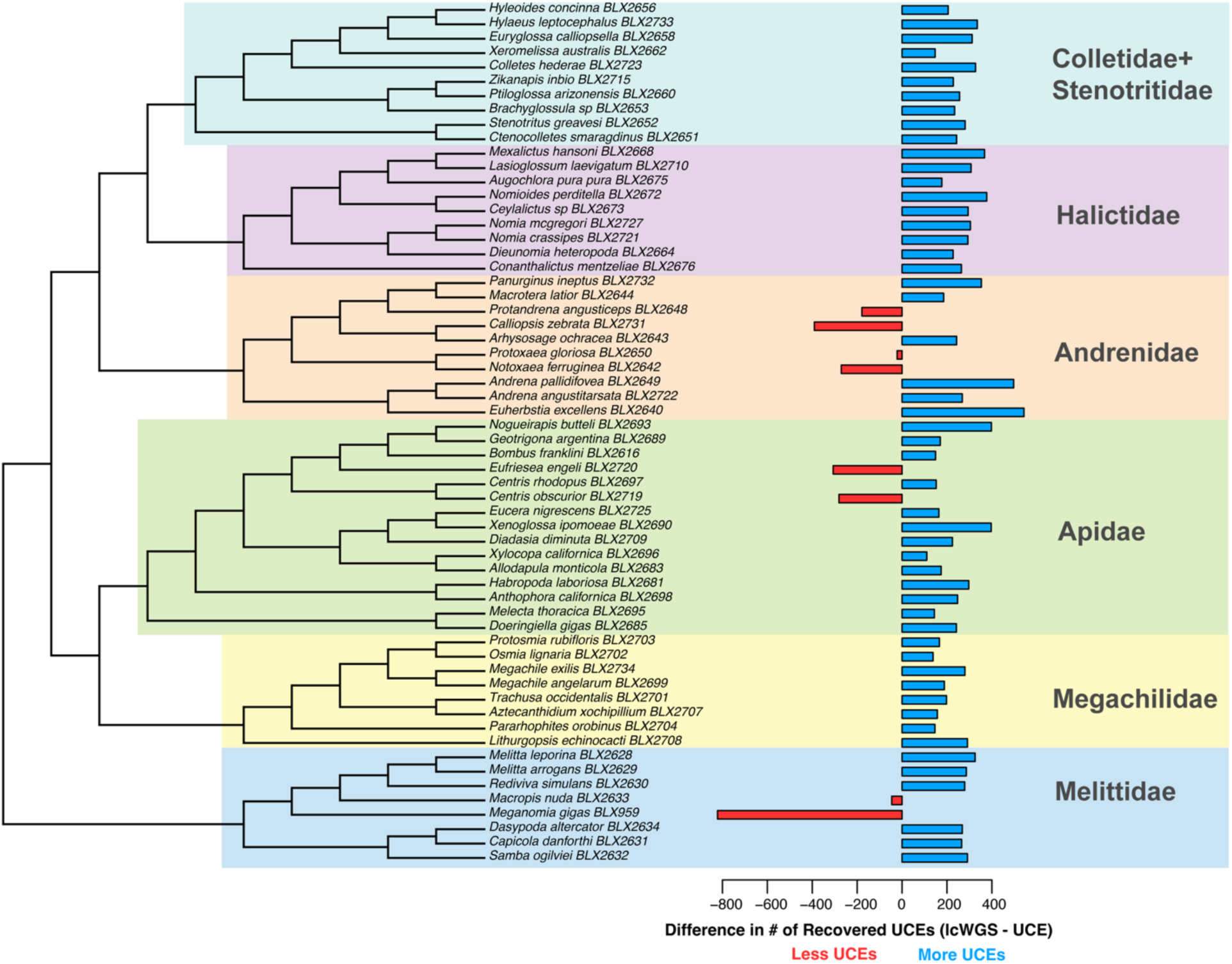
Phylogenetic distribution of UCE recovery differences between lcWGS and UCE-enriched datasets. Tree topology inferred from the concatenated lcWGS dataset using IQ-TREE v2 (GTR+F+G4). All nodes received 100% ultrafast bootstrap (UFB) support. For each taxon, bars indicate relative UCE locus recovery: blue = more loci recovered with lcWGS; red = more loci recovered with UCE enrichment. Only 8 of 60 samples showed fewer loci recovered under lcWGS.

### Phylogenetic Accuracy

Concatenated phylogenetic analyses of the lcWGS and UCE-enriched datasets yielded largely congruent topologies, with all nodes receiving maximal UFB support. The only topological discrepancy involved the placement of *Brachyglossula* sp. (BLX2653; Neopasiphaeinae), which was recovered as sister to *Ptiloglossa* + *Zikanapis* in the lcWGS phylogeny, but sister to all other colletid bees in the UCE-enriched phylogeny (Fig. 4; Fig. S2 and S3).

When focusing on individual gene trees, lcWGS consistently showed improved performance. Mean UFB support was significantly higher in lcWGS-derived gene trees. Gene trees from lcWGS were also significantly more similar to the reference topology based on quartet similarity scores. Gene concordance factors (gCF) were higher for lcWGS under both reference trees, although these differences were not statistically significant (Figs. 5 and S4).

**Figure 5.**
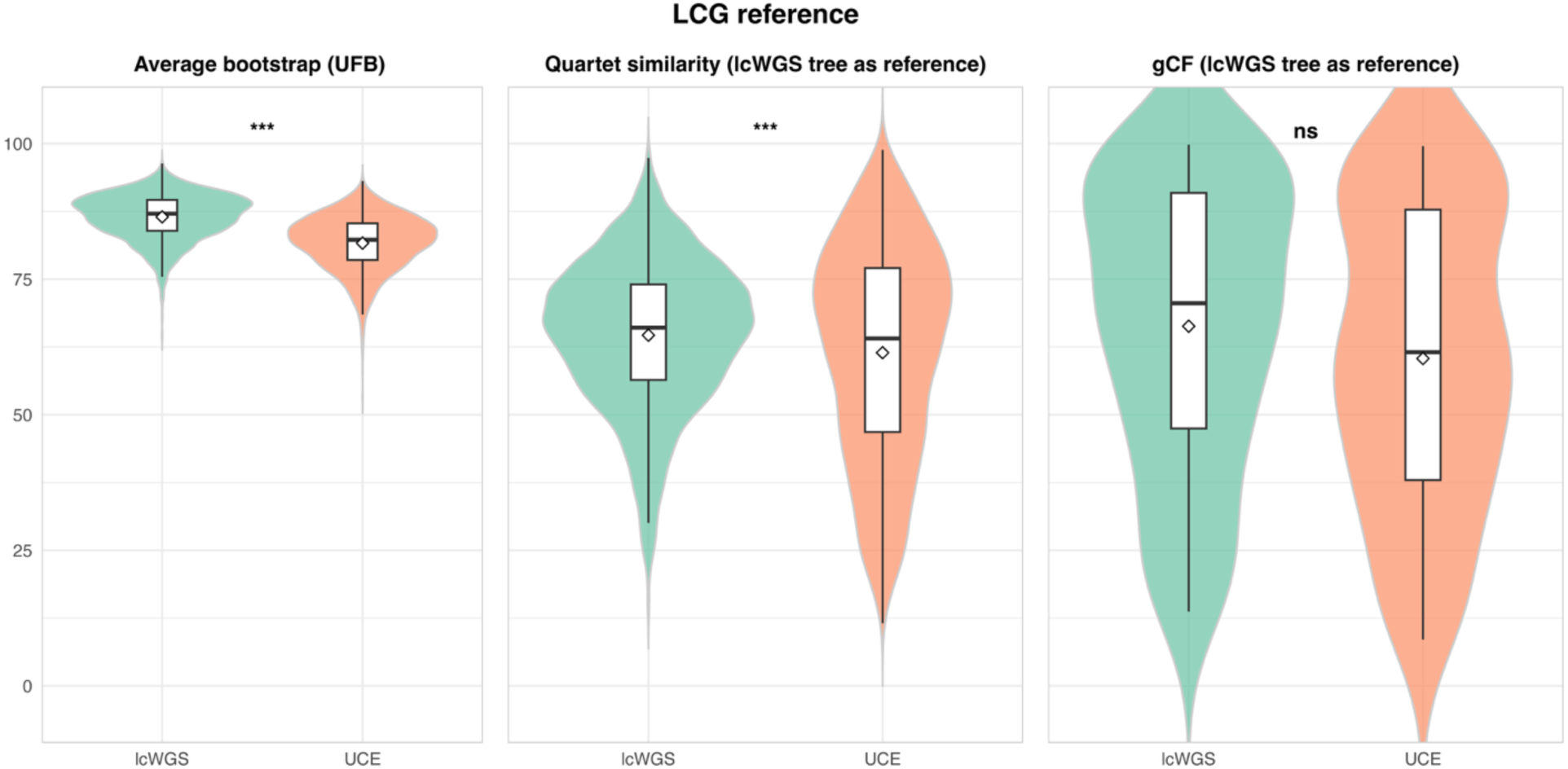
Comparison of gene tree performance metrics between lcWGS and UCE-enriched datasets. Violin and box plots show (left to right): mean ultrafast bootstrap (UFB) support across gene trees, quartet similarity to the reference topology (gene tree “accuracy”), and gene concordance factors (gCF) across reference tree nodes (n = 57 nodes). Statistical significance was assessed using Wilcoxon signed-rank tests with multiple-testing correction where applicable. Asterisks denote significance levels (*** p < 0.0001; ns = not significant).

### Recovery of BUSCOs, Mitogenomes, and Apoidea UCEs

The lcWGS dataset substantially outperformed the UCE-enriched dataset in the recovery of additional marker types, including Hymenoptera BUSCO genes, mitochondrial genes, and loci from the expanded Apoidea UCE probe set (Figs. 6, S8, and S9; Table 2 and S7). Across all samples, lcWGS recovered significantly more Hymenoptera_odb10 BUSCO genes (mean: 3,538 vs. 155; p < 0.001), mitochondrial genes (13 vs. 8; p < 0.001), and Apoidea UCE loci (12,473 vs. 5,014; p < 0.001). When stratified by age category, lcWGS outperformed UCE enrichment for all metrics in all age groups. All comparisons were statistically significant, excluding mitochondrial gene recovery in the most recent age category. Similar results were observed for other BUSCO gene sets (Fig. S8) and for additional Apoidea UCE metrics (Fig. S9).

**Figure 6.**
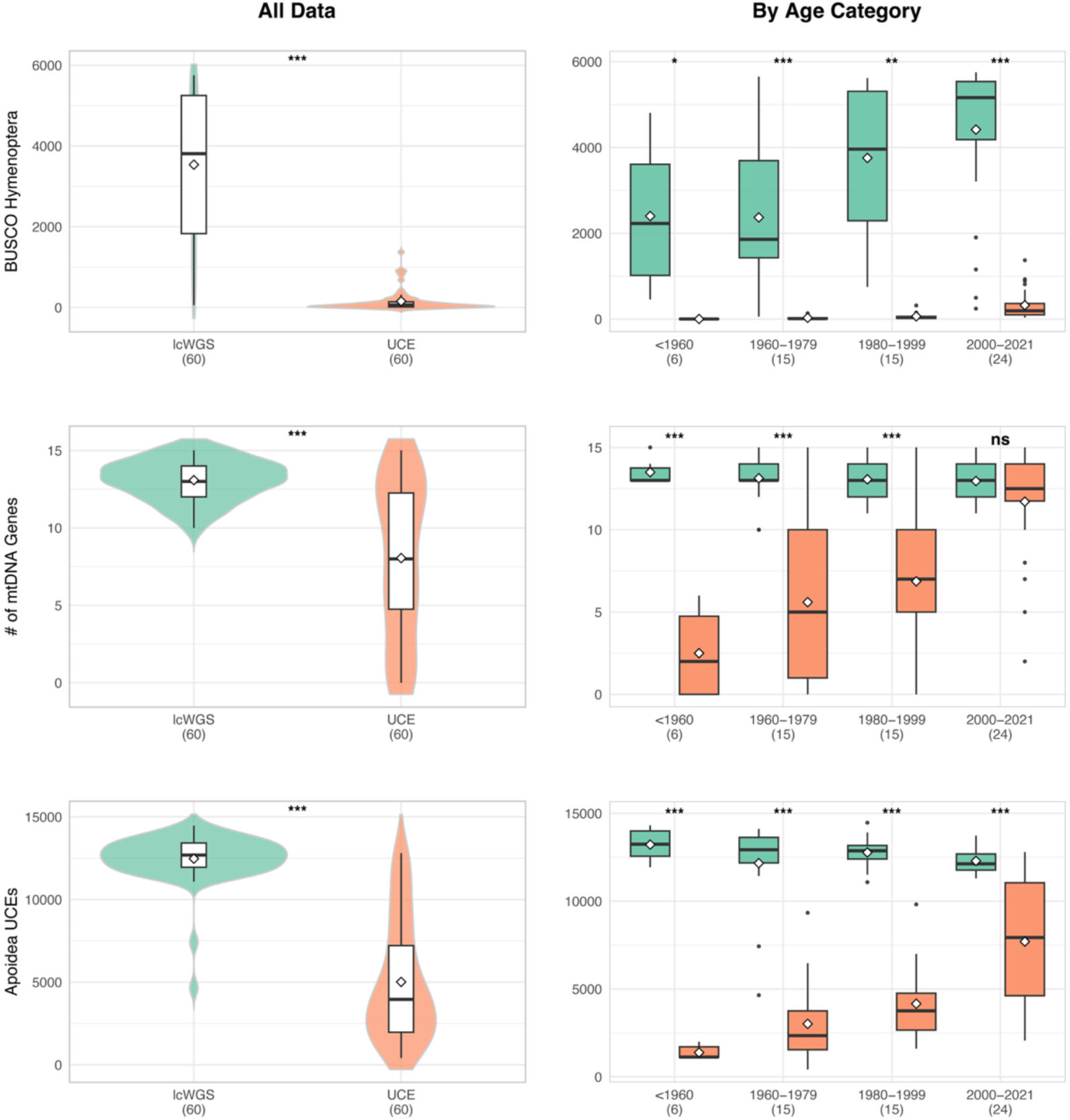
Recovery of additional marker types from lcWGS and UCE-enriched datasets. Left column shows all samples combined; right column shows results by age category. From top to bottom: (1) number of Hymenoptera_odb10 BUSCO genes recovered, (2) number of mitochondrial genes identified, and (3) number of Apoidea 20Kv1 UCE loci extracted *in silico*. Violin and box plots summarize distributions. Paired statistical comparisons between lcWGS and UCE datasets were performed per sample (Wilcoxon signed-rank or paired t-test as appropriate; BH-adjusted p-values). Asterisks indicate significance levels. All comparisons are paired by sample.

In the UCE-enriched dataset, all three marker classes showed significant positive correlations with collection year (Fig. 7), indicating reduced performance in older specimens. In contrast, lcWGS showed a significant positive correlation with collection year only for BUSCO gene recovery. The other lcWGS marker types exhibited weak, non-significant negative correlations with age.

**Figure 7.**
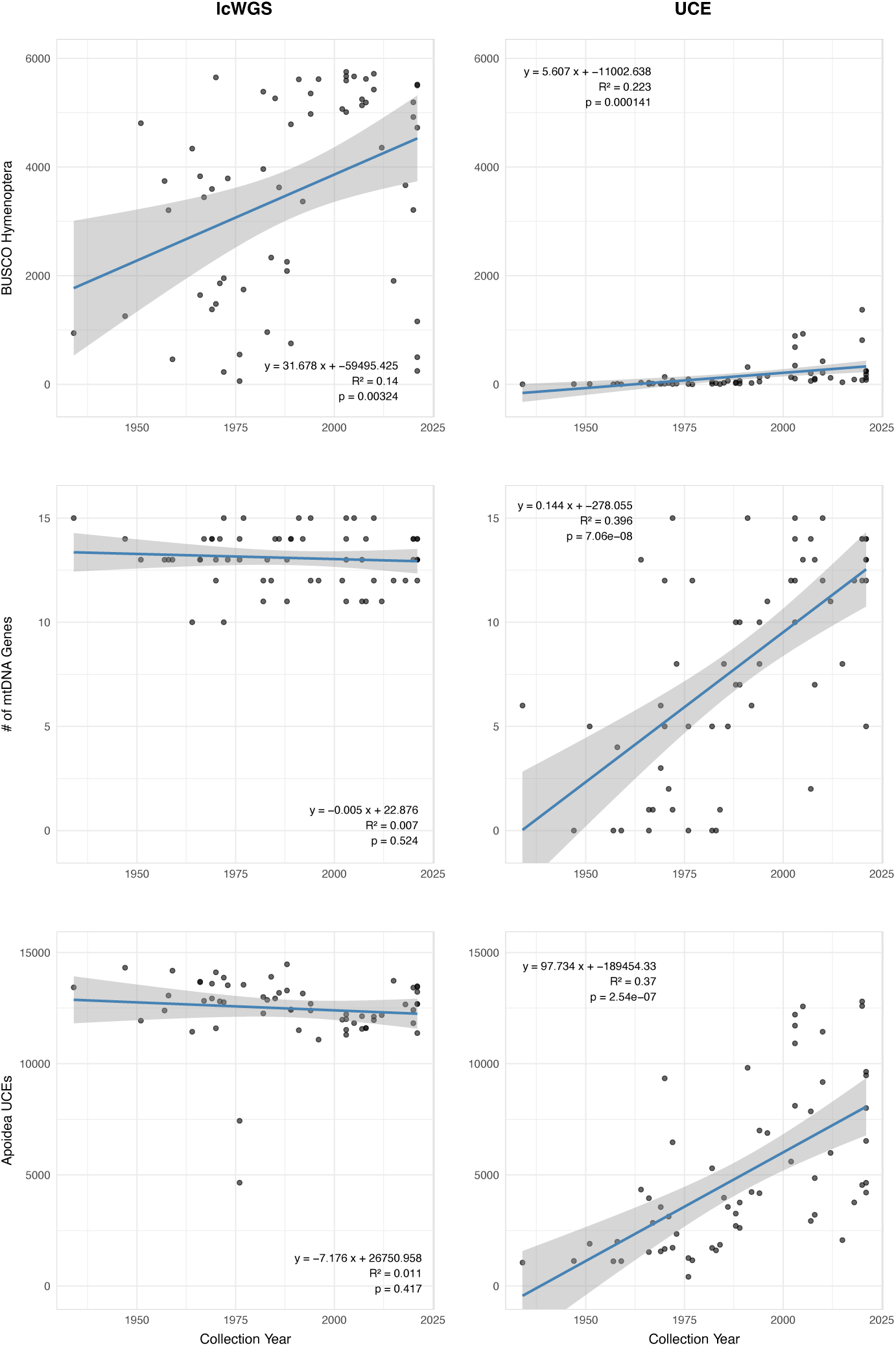
Relationships between recovery of additional marker types and specimen collection year. Left column shows lcWGS results; right column shows UCE-enriched results. From top to bottom: number of Hymenoptera_odb10 BUSCO genes, mitochondrial genes, and Apoidea 20Kv1 UCE loci recovered per sample. Solid lines represent ordinary least squares regression fits. Regression statistics are provided in Table 2.

## Discussion

### Ultraconserved element data quality – lcWGS vs. UCE enrichment

The rapid expansion of phylogenomics has reshaped systematic biology, yet the field remains anchored to reduced-representation methods developed when sequencing costs were substantially higher (Faircloth et al. 2012, Lemmon and Lemmon 2013, McCormack et al. 2013, Andermann et al. 2020, Young and Gillung 2020). As whole-genome sequencing becomes increasingly affordable, it is critical to reassess whether targeted enrichment remains the optimal strategy—particularly for museum specimens, which represent an irreplaceable archive of biodiversity (Suchan et al. 2016, Raxworthy and Smith 2021). Despite use of both hybrid capture and genome skimming approaches, rigorous head-to-head comparisons under realistic museomic conditions are lacking. By generating lcWGS and UCE-enriched datasets from identical Illumina libraries spanning nearly a century of bee specimens, we provide a controlled benchmark of methodological performance. Our results show that lcWGS not only matches but frequently exceeds UCE enrichment in locus yield, locus length, and phylogenetic performance, with the greatest advantages observed in older, more degraded specimens.

Across the full dataset and within age categories, lcWGS recovered more UCE loci and substantially longer loci than UCE enrichment (Fig. 2). Although both approaches showed some decline in performance with increasing specimen age, this effect was consistently stronger in the enrichment dataset (Fig. 3), indicating greater sensitivity to DNA degradation. These findings suggest that hybrid capture, while powerful, introduces a recovery bottleneck that becomes increasingly restrictive as fragment length and DNA integrity decline.

The mechanistic explanation for this difference is straightforward. Target enrichment relies on successful hybridization between library fragments and probe sequences. Only fragments that bind probes are retained at high coverage, and flanking sequence is recovered only if it resides on the same fragment as the probe-binding region (Bi et al. 2013, Tin et al. 2014, Zhang, Y. Miles et al. 2019b, Andermann et al. 2020, Freitas et al. 2023). In degraded specimens, shorter fragment sizes and chemical damage reduce the likelihood of efficient hybridization and limit the recovery of extended flanking regions (Paijmans et al. 2016, Suchan et al. 2016). By contrast, lcWGS eliminates this hybridization bottleneck. All DNA fragments have an equal probability of being sequenced, and overlapping reads can be assembled into longer contiguous regions without requiring physical linkage to probe targets. This difference in recovery architecture explains the consistent superiority of lcWGS across specimen ages.

Importantly, lcWGS recovered fewer UCE loci in 8 of 60 samples. These exceptions likely reflect genome size variation among taxa (Gregory 2005, Gregory et al. 2007, Cong et al. 2022) and non-target sequence contamination. Because lcWGS distributes sequencing effort across the entire genome, larger genomes result in reduced effective per-locus coverage. In taxa with genomes exceeding the assumed ∼500 Mb baseline used for coverage estimation, realized depth at UCE loci may have been lower than optimal. In addition, historical museum specimens are susceptible to post-mortem DNA degradation and colonization by environmental microbes, including fungi and bacteria, which can contribute substantial amounts of exogenous DNA (Staats et al. 2013). In low-coverage sequencing libraries, such contamination can further dilute endogenous signal and reduce recovery of target loci. Consistent with this expectation, assembly diagnostics for several low-performing samples indicated elevated levels of non-target sequence content and reduced assembly completeness, supporting contamination as a contributing factor. In contrast, enrichment concentrates sequencing effort on predefined targets regardless of genome size and may partially mitigate the effects of both genome-scale dilution and exogenous DNA. Thus, genome size remains a meaningful predictor of lcWGS efficiency, while occasional reversals in performance likely reflect a combination of biological variation and specimen-specific factors such as contamination. Incorporating genome size estimates and assessing contamination levels during study design may further improve locus recovery and sequencing efficiency.

### Implications for phylogenetic accuracy and gene-tree resolution

While concatenated trees from both datasets were largely congruent and maximally supported, analyses of individual gene trees reveal more consequential differences. lcWGS-derived loci exhibited significantly higher mean bootstrap support and greater quartet similarity to reference topologies (Fig. 5), indicating reduced gene-tree estimation error. Gene concordance factors were also consistently higher for lcWGS, although these differences were not statistically significant.

Gene-tree estimation error is a persistent challenge in phylogenomics and can be misinterpreted as biological signal, including incomplete lineage sorting or introgression (Edwards et al. 2016, Springer and Gatesy 2017). Simulation and empirical studies demonstrate that shorter loci are particularly susceptible to stochastic error and model misspecification (Sayyari et al. 2017). By increasing locus length and per-locus informativeness, lcWGS appears to reduce stochastic error and improve gene-tree stability. This effect is particularly important when degraded museum specimens are included, as short, information-poor loci are more susceptible to erroneous resolution (Freitas et al. 2023). Even modest increases in locus length, or information content, can substantially improve species-tree inference when large numbers of loci are combined (Sayyari and Mirarab 2016). Thus, improvements in locus recovery translate directly into improved phylogenetic robustness.

Another critical distinction is flexibility. Low-coverage WGS allows flanking regions of UCEs to be extended *in silico* without additional laboratory investment, permitting researchers to tailor locus length and informativeness to the evolutionary timescale of interest. Increasing probe tiling density in enrichment experiments may extend target recovery to some extent, but this increases cost and remains constrained by fragment hybridization. In contrast, lcWGS decouples locus definition from laboratory design, enabling dynamic redefinition of marker boundaries as analytical goals evolve.

### Broad genomic representation and reusability of lcWGS data

Beyond UCE recovery, lcWGS also enabled recovery of a wider diversity of genomic markers (Fig. 6). BUSCO gene recovery was an order of magnitude higher in lcWGS assemblies, nearly complete mitochondrial gene sets were recovered for most samples, and use of the expanded Apoidea bait set yielded several-fold more UCE loci. Notably, >90% of BUSCO genes were recovered in 12 of the 60 lcWGS samples, indicating that relatively complete gene space can be obtained even from degraded museum specimens. This level of completeness suggests that near-comprehensive sets of protein-coding genes, including much of the exome, are likely recoverable for a substantial fraction of samples, enabling downstream comparative genomic analyses beyond phylogenetics, including gene family evolution, molecular adaptation, and trait-associated genomic variation. While these results were intuitively expected based on the nature of the methods, empirical demonstration remains important because target enrichment is not a perfect process, with off-target reads from the mitogenome and broader nuclear genome expected to some degree (Raposo do Amaral et al. 2015, Allio et al. 2020, Andermann et al. 2020, Swami et al. 2026).

Our results demonstrate that lcWGS provides a highly flexible substrate for multi-marker discovery. The same datasets can be mined for diverse genomic targets, including organellar genomes, single-copy nuclear genes, and probe-defined loci, enabling iterative reuse as analytical methods and research questions evolve. In contrast, enrichment datasets are inherently limited to the targeted loci and are less easily repurposed once generated. The capacity of lcWGS data to support multiple marker systems—from organellar to nuclear single-copy genes—offers clear advantages for large-scale comparative studies, biodiversity inventories, and future meta-analyses that seek to integrate disparate datasets.

Genome skimming approaches have long been used to recover organellar genomes and high-copy markers (Zeng et al. 2018, Olofsson et al. 2019, Liu et al. 2021, Taite et al. 2023, Hu et al. 2024), and shallow WGS has been shown to recover single-copy nuclear loci suitable for phylogenomics (Zhang, Feng et al. 2019a). Our results extend these findings to degraded insect museum specimens and demonstrate that lcWGS can outperform targeted enrichment even for loci originally designed for in-solution capture.

### Practical considerations for lcWGS implementation

Despite its performance advantages, the utility of lcWGS is influenced by genome size, sequencing cost, and computational resources. As shown above, lcWGS not only recovered more loci across multiple marker types but also provided broader genomic representation, increasing its value per dataset relative to targeted enrichment. For small-genome taxa such as bees (approximately 200–800 Mb) (Alfsnes et al. 2017, Sproul et al. 2023, Cook et al. 2025), sequencing to ∼20× coverage per specimen is increasingly feasible at scale and can approach cost parity with targeted enrichment when probe synthesis, hybridization, and labor are considered (Zhang, Y. Miles et al. 2019b). Assuming a 500 Mb genome, an Illumina NovaSeq 25B lane (2 × 150 bp) generating ∼1 Tb of sequence data could theoretically accommodate ∼100 samples at 20× coverage. At an approximate cost of $2,700 per lane, this corresponds to a sequencing cost on the order of ∼$27 per sample, although realized costs will vary depending on data yield, multiplexing efficiency, and library performance. In practice, the number of samples per lane can be adjusted based on genome size, DNA quality, and target coverage.

For taxa with larger or more variable genomes, targeted enrichment may remain more economical until sequencing costs decline further. Low-coverage WGS also requires greater computational storage and processing time; however, these demands are offset by streamlined laboratory workflows and the elimination of enrichment steps, which can reduce failure rates and inter-batch variation. The simplification of molecular workflows may be particularly advantageous for large-scale projects or institutions processing many specimens of uncertain DNA quality.

### Limitations and future directions

This study was limited to short-read Illumina data, which can be complemented by long-read lcWGS to further improve assembly contiguity and recover repetitive or GC-rich regions. Our sampling focused on bees, and extending these analyses to other organismal groups will be important for evaluating generality across taxa with different genome sizes and architectures. Future work could also explore hybrid strategies that combine shallow lcWGS with moderate enrichment, thereby leveraging the advantages of both approaches. Additional studies should investigate optimal sequencing coverage thresholds for phylogenomic applications, including how assembly quality, locus recovery, and downstream phylogenetic inference change across varying coverage levels. It will also be important to evaluate optimal flank lengths for extracting UCE regions from lcWGS assemblies, as longer flanking regions may provide substantially greater phylogenetic signal while also increasing challenges related to alignment and homology assessment. Finally, development of standardized workflows for extracting homologous loci from lcWGS and enrichment datasets would facilitate integration of legacy and newly generated data within unified phylogenomic frameworks.

## Conclusions

Low-coverage whole-genome sequencing provides a robust and broadly informative alternative to target enrichment for phylogenomics, particularly for museum specimens with degraded DNA. Compared to UCE enrichment, lcWGS yields more loci, longer alignments, higher gene-tree support, and data that can be readily repurposed for diverse downstream analyses. When combined with increasing cost-efficiency and scalable sequencing strategies, these advantages enhance the overall value of lcWGS datasets relative to enrichment approaches. As sequencing costs continue to decline, lcWGS is likely to replace targeted enrichment for many small-genome taxa, enabling a unified framework that bridges phylogenetics, comparative genomics, population genomics, and museomics from a single, reusable genomic resource.

## Funding

This work was supported by the U.S. National Science Foundation (grant numbers DEB-2127744, DEB-2127745); U.S. Department of Agriculture (in-house project number 2080-30500-001-000D); Coordenação de Aperfeiçoamento de Pessoal de Nível Superior (finance code 001); and São Paulo Research Foundation (grant numbers 2023/04048-0, 2024/12303-2).

## Acknowledgements

We thank Kerrigan Tobin for laboratory work. We thank Terry Griswold and Harold Ikerd from the U.S. National Pollinating Insects Collection for help with specimen databasing and curation. This research used computing resources provided by the USDA-ARS SCINet project and/or the AI Center of Excellence (0201-88888-003-000D; 0201-88888-002-000D). USDA is an equal opportunity provider and employer.

## Data Availability

All data and analysis materials necessary to reproduce the analyses are archived on Dryad (DOI: https://doi.org/10.5061/dryad.fbg79cpb8. This archive includes SPAdes assemblies, Hymenoptera and Apoidea UCE contigs, phylogenetic matrices, concatenated and gene-tree files, analysis inputs, scripts, statistical outputs, and generated figures. The version-controlled analysis repository is available on GitHub at https://github.com/branstetter-lab/lcg_uce_paper. Raw sequencing reads have been deposited in the NCBI Sequence Read Archive under BioProject accession PRJNA1261093.

## Supplemental Materials

### Supplemental Figures

Figure S1. Reads vs. specimen age.

Figure S2. Trimmed reads vs. specimen age.

Figure S3. GenomeScope2 genome size estimates.

Figure S4. Sequencing coverage for bulk and UCE contigs.

Figure S5. Concatenated phylogeny based on the lcWGS enriched loci.

Figure S6. Concatenated phylogeny based on the UCE enriched loci.

Figure S7. Gene tree performance based on the UCE concatenated reference topology.

Figures S8. Complete BUSCO gene recovery results.

Figures S9. Apoidea UCE locus recovery and length results.

Figure S10. Concatenated phylogeny based on Apoidea UCEs extracted from the lcWGS data.

### Supplemental Tables

Table S1. Sample voucher data.

Table S2. Sample laboratory data.

Table S3. NCBI accession numbers.

Table S4. Genome sample data for Apoidea probe set development.

Table S5. Sequencing, assembly, and UCE stats for lcWGS and UCE datasets.

Table S6. Statistical analyses comparing sample age and read counts.

Table S7. BUSCO, mtDNA, and Apoidea UCE stats for lcWGS and UCE datasets.

## Notes

### Competing Interest Statement

The authors have declared no competing interest.

## References

Alfsnes K., Leinaas H.P., Hessen D.O. 2017. Genome size in arthropods; different roles of phylogeny, habitat and life history in insects and crustaceans. Ecology and Evolution 7:5939–5947.

Allen J.M., Boyd B., Nguyen N.P., Vachaspati P., Warnow T., Huang D.I., Grady P.G.S., Bell K.C., Cronk Q.C.B., Mugisha L., Pittendrigh B.R., Leonardi M.S., Reed D.L., Johnson K.P. 2017. Phylogenomics from whole genome sequences using aTRAM. Syst. Biol. 66:786–798.

Allio R., Schomaker-Bastos A., Romiguier J., Prosdocimi F., Nabholz B., Delsuc F. 2020. MitoFinder: efficient automated large-scale extraction of mitogenomic data in target enrichment phylogenomics. Molecular Ecology Resources.

Almeida E.A., Bossert S., Danforth B.N., Porto D.S., Freitas F.V., Davis C.C., Murray E.A., Blaimer B.B., Spasojevic T., Ströher P.R., Orr M.C., Packer L., Brady S.G., Kuhlman M., Branstetter M.G., Pie M.R. 2023. The evolutionary history of bees in time and space. Curr. Biol. 33:3409–3422.

Andermann T., Torres Jimenez M.F., Matos-Maravi P., Batista R., Blanco-Pastor J.L., Gustafsson A.L.S., Kistler L., Liberal I.M., Oxelman B., Bacon C.D., Antonelli A. 2020. A guide to carrying out a phylogenomic target sequence capture project. Frontiers in Genetics 10:1407.

Andrews S. 2010. FastQC: A quality control tool for high throughput sequence data.

Asher R.J., Smith M.R., Davalos L. 2022. Phylogenetic signal and bias in paleontology. Syst. Biol. 71:986–1008.

Baca S.M., Toussaint E.F.A., Miller K.B., Short A.E.Z. 2017. Molecular phylogeny of the aquatic beetle family Noteridae (Coleoptera: Adephaga) with an emphasis on data partitioning strategies. Mol. Phylogen. Evol. 107:282–292.

Bankevich A., Nurk S., Antipov D., Gurevich A.A., Dvorkin M., Kulikov A.S., Lesin V.M., Nikolenko S.I., Pham S., Prjibelski A.D., Pyshkin A.V., Sirotkin A.V., Vyahhi N., Tesler G., Alekseyev M.A., Pevzner P.A. 2012. SPAdes: a new genome assembly algorithm and its applications to single-cell sequencing. J. Comput. Biol. 19:455–477.

Bi K., Linderoth T., Vanderpool D., Good J.M., Nielsen R., Moritz C. 2013. Unlocking the vault: next-generation museum population genomics. Mol. Ecol. 22:6018–6032.

Blaimer B.B., Lloyd M.W., Guillory W.X., Brady S.G. 2016. Sequence capture and phylogenetic utility of genomic Ultraconserved Elements obtained from pinned insect specimens. Plos One 11.

Branstetter M.G., Danforth B.N., Pitts J.P., Faircloth B.C., Ward P.S., Buffington M.L., Gates M.W., Kula R.R., Brady S.G. 2017a. Phylogenomic insights into the evolution of stinging wasps and the origins of ants and bees. Curr. Biol. 27:1019–1025.

Branstetter M.G., Longino J.T. 2019. Ultra-conserved element phylogenomics of new world *Ponera* (Hymenoptera: Formicidae) illuminates the origin and phylogeographic history of the endemic exotic ant *Ponera exotica*. Insect Systematics and Diversity 3:1-13.

Branstetter M.G., Longino J.T., Ward P.S., Faircloth B.C. 2017b. Enriching the ant tree of life: enhanced UCE bait set for genome-scale phylogenetics of ants and other Hymenoptera. Methods in Ecology and Evolution 8:768–776.

Bushnell B. 2014. BBTools software package. URL http://sourceforge.net/projects/bbmap.

Chen K.-Y., Wang J.-D., Xiang R.-Q., Yang X.-D., Yun Q.-Z., Huang Y., Sun H., Chen J.-H. 2025. Backbone phylogeny of *Salix* based on genome skimming data. Plant Diversity 47:178–188.

Cong Y., Ye X., Mei Y., He K., Li F. 2022. Transposons and non-coding regions drive the intrafamily differences of genome size in insects. iScience 25:104873.

Cook H.L., Sproul J.S., Murray E.A., Bossert S. 2025. A comparative analysis of transposable element diversity and evolution across 75 bee genomes. BMC Genomics 26.

Edwards S.V., Xi Z., Janke A., Faircloth B.C., McCormack J.E., Glenn T.C., Zhong B., Wu S., Lemmon E.M., Lemmon A.R., Leache A.D., Liu L., Davis C.C. 2016. Implementing and testing the multispecies coalescent model: A valuable paradigm for phylogenomics. Mol Phylogenet Evol 94:447–462.

Ewels P., Magnusson M., Lundin S., Kaller M. 2016. MultiQC: summarize analysis results for multiple tools and samples in a single report. Bioinformatics 32:3047–3048.

Faircloth B.C. 2013. illumiprocessor: a trimmomatic wrapper for parallel adapter and quality trimming.

Faircloth B.C. 2016. PHYLUCE is a software package for the analysis of conserved genomic loci. Bioinformatics 32:786–788.

Faircloth B.C., McCormack J.E., Crawford N.G., Harvey M.G., Brumfield R.T., Glenn T.C. 2012. Ultraconserved elements anchor thousands of genetic markers spanning multiple evolutionary timescales. Syst. Biol. 61:717–726.

Freitas F.V., Branstetter M.G., Franceschini-Santos V.H., Dorchin A., Wright K.W., López-Uribe M.M., Griswold T., Silveira F.A., Almeida E.A.B., Gillung J. 2023. UCE phylogenomics, biogeography, and classification of long-horned bees (Hymenoptera: Apidae: Eucerini), with insights on using specimens with extremely degraded DNA. Insect Systematics and Diversity 7.

Gillung J.P., Winterton S.L., Bayless K.M., Khouri Z., Borowiec M.L., Yeates D., Kimsey L.S., Misof B., Shin S., Zhou X., Mayer C., Petersen M., Wiegmann B.M. 2018. Anchored phylogenomics unravels the evolution of spider flies (Diptera, Acroceridae) and reveals discordance between nucleotides and amino acids. Mol Phylogenet Evol 128:233–245.

Glenn T.C., Nilsen R.A., Kieran T.J., Sanders J.G., Bayona-Vásquez N.J., Finger J.W., Pierson T.W., Bentley K.E., Hoffberg S.L., Louha S., Garcia-De Leon F.J., Del Rio Portilla M.A., Reed K.D., Anderson J.L., Meece J.K., Aggrey S.E., Rekaya R., Alabady M., Belanger M., Winker K., Faircloth B.C. 2019. Adapterama I: universal stubs and primers for 384 unique dual-indexed or 147,456 combinatorially-indexed Illumina libraries (iTru & iNext). PeerJ 7:e7755.

Goodman A., Tolman E., Uche-Dike R., Abbott J., Breinholt J.W., Bybee S., Frandsen P.B., Gosnell J.S., Guralnick R., Kalkman V.J., Kohli M., Lontchi J.F., Lupiyaningdyah P., Newton L., Ware J.L., Yoshizawa K. 2023. Assessment of targeted enrichment locus capture across time and museums using odonate specimens. Insect Systematics and Diversity 7.

Grab H., Branstetter M.G., Amon N., Urban-Mead K.R., Park M.G., Gibbs J., Blitzer E.J., Poveda K., Loeb G., Danforth B.N. 2019. Agriculturally dominated landscapes reduce bee phylogenetic diversity and pollination services. Science 363:282–284.

Gregory T.R. 2005. The Evolution of the Genome. El Sevier, p. 731.

Gregory T.R., Nicol J.A., Tamm H., Kullman B., Kullman K., Leitch I.J., Murray B.G., Kapraun D.F., Greilhuber J., Bennett M.D. 2007. Eukaryotic genome size databases. Nucleic Acids Res. 35:D332–338.

Harris R.S. 2007. Improved pairwise alignment of genomic DNA. Pennsylvania State University.

Hedin M., Derkarabetian S., Alfaro A., Ramirez M.J., Bond J.E. 2019. Phylogenomic analysis and revised classification of atypoid mygalomorph spiders (Araneae, Mygalomorphae), with notes on arachnid ultraconserved element loci. PeerJ 7:e6864.

Hoang D.T., Chernomor O., von Haeseler A., Minh B.Q., Vinh L.S. 2018. UFBoot2: improving the ultrafast bootstrap approximation. Mol. Biol. Evol. 35:518–522.

Hu X.-Z., Guo C., Qin S.-Y., Li D.-Z., Guo Z.-H. 2024. Deep genome skimming reveals the hybrid origin of *Pseudosasa gracilis* (Poaceae: Bambusoideae). Plant Diversity 46:344–352.

Huang W., Li L., Myers J.R., Marth G.T. 2012. ART: a next-generation sequencing read simulator. Bioinformatics 28:593–594.

Katoh K., Standley D.M. 2013. MAFFT multiple sequence alignment software version 7: improvements in performance and usability. Mol. Biol. Evol. 30:772–780.

Kent W.J. 2002. BLAT—The BLAST-like alignment tool. Genome Res. 12:656–664.

Laetsch D.R., Blaxter M.L. 2017. BlobTools: Interrogation of genome assemblies. F1000Research 6.

Lemmon E.M., Lemmon A.R. 2013. High-throughput genomic data in systematics and phylogenetics. Annual Review of Ecology, Evolution, and Systematics 44:99–121.

Liu B.B., Ma Z.Y., Ren C., Hodel R.G.J., Sun M., Liu X.Q., Liu G.N., Hong D.Y., Zimmer E.A., Wen J. 2021. Capturing single-copy nuclear genes, organellar genomes, and nuclear ribosomal DNA from deep genome skimming data for plant phylogenetics: A case study in Vitaceae. Journal of Systematics and Evolution 59:1124–1138.

Manni M., Berkeley M.R., Seppey M., Simão F.A., Zdobnov E.M., Kelley J. 2021. BUSCO update: Novel and streamlined workflows along with broader and deeper phylogenetic coverage for scoring of eukaryotic, prokaryotic, and viral genomes. Mol. Biol. Evol. 38:4647–4654.

McCormack J.E., Hird S.M., Zellmer A.J., Carstens B.C., Brumfield R.T. 2013. Applications of next-generation sequencing to phylogeography and phylogenetics. Mol. Phylogen. Evol. 66:526–538.

Mikheyev A.S., Zwick A., Magrath M.J.L., Grau M.L., Qiu L., Su Y.N., Yeates D. 2017. Museum genomics confirms that the Lord Howe Island stick insect survived extinction. Curr. Biol. 27:3157–3161 e3154.

Minh B.Q., Hahn M.W., Lanfear R., Rosenberg M. 2020a. New methods to calculate concordance factors for phylogenomic datasets. Mol. Biol. Evol. 37:2727–2733.

Minh B.Q., Nguyen M.A.T., von Haeseler A. 2013. Ultrafast approximation for phylogenetic bootstrap. Mol. Biol. Evol. 30:1188–1195.

Minh B.Q., Schmidt H.A., Chernomor O., Schrempf D., Woodhams M.D., von Haeseler A., Lanfear R. 2020b. IQ-TREE 2: new models and efficient methods for phylogenetic inference in the genomic era. Mol. Biol. Evol. 37:1530–1534.

Olofsson J.K., Cantera I., Van de Paer C., Hong-Wa C., Zedane L., Dunning L.T., Alberti A., Christin P.A., Besnard G. 2019. Phylogenomics using low-depth whole genome sequencing: A case study with the olive tribe. Molecular Ecology Resources 19:877–892.

Paijmans J.L., Fickel J., Courtiol A., Hofreiter M., Forster D.W. 2016. Impact of enrichment conditions on cross-species capture of fresh and degraded DNA. Molecular Ecology Resources 16:42–55.

Pezzini F.F., Ferrari G., Forrest L.L., Hart M.L., Nishii K., Kidner C.A. 2023. Target capture and genome skimming for plant diversity studies. Applications in Plant Sciences 11.

Ranallo-Benavidez T.R., Jaron K.S., Schatz M.C. 2020. GenomeScope 2.0 and Smudgeplot for reference-free profiling of polyploid genomes. Nature Communications 11:1432.

Raposo do Amaral F., Neves L.G., Resende M.F., Jr., Mobili F., Miyaki C.Y., Pellegrino K.C., Biondo C. 2015. Ultraconserved elements sequencing as a low-cost source of complete mitochondrial genomes and microsatellite markers in non-model amniotes. PLoS One 10:e0138446.

Raxworthy C.J., Smith B.T. 2021. Mining museums for historical DNA: Advances and challenges in museomics. Trends Ecol. Evol. 36:1049–1060.

Rohland N., Reich D. 2012. Cost-effective, high-throughput DNA sequencing libraries for multiplexed target capture. Genome Res. 22:939–946.

Sayyari E., Mirarab S. 2016. Fast Coalescent-Based Computation of Local Branch Support from Quartet Frequencies. Mol. Biol. Evol. 33:1654–1668.

Sayyari E., Whitfield J.B., Mirarab S. 2017. Fragmentary Gene Sequences Negatively Impact Gene Tree and Species Tree Reconstruction. Mol. Biol. Evol. 34:3279–3291.

Schweizer R.M., Grummer J.A., Tobin K.B., Corpuz R., Geib S.M., Cox-Foster D., Kimsey L.S., Koch J.B.U., Branstetter M.G. 2025. Museum genomics suggests long-term population decline in a putatively extinct bumble bee. Proceedings of the National Academy of Sciences 43:e2509749122.

Smith M.R. 2019. Quartet: comparison of phylogenetic trees using quartet and split measures. R package version 1.2.

Springer M.S., Gatesy J. 2017. On the importance of homology in the age of phylogenomics. Syst. Biodivers.:1–19.

Sproul J.S., Hotaling S., Heckenhauer J., Powell A., Marshall D., Larracuente A.M., Kelley J.L., Pauls S.U., Frandsen P.B. 2023. Analyses of 600+ insect genomes reveal repetitive element dynamics and highlight biodiversity-scale repeat annotation challenges. Genome Res. 33:1708–1717.

Staats M., Erkens R.H., van de Vossenberg B., Wieringa J.J., Kraaijeveld K., Stielow B., Geml J., Richardson J.E., Bakker F.T. 2013. Genomic treasure troves: complete genome sequencing of herbarium and insect museum specimens. PLoS One 8:e69189.

Suchan T., Pitteloud C., Gerasimova N.S., Kostikova A., Schmid S., Arrigo N., Pajkovic M., Ronikier M., Alvarez N. 2016. Hybridization capture using RAD probes (hyRAD), a new tool for performing genomic analyses on collection specimens. PLoS ONE 11:e0151651.

Swami A.V., Baraf L.M., Cowman P.F. 2026. Mining for mitochondria: 68 mitogenomes for wrasses and parrotfishes (F: Labridae) from off-target UCE data. Ecology and Evolution 16.

Taite M., Fernández-Álvarez F.Á., Braid H.E., Bush S.L., Bolstad K., Drewery J., Mills S., Strugnell J.M., Vecchione M., Villanueva R., Voight J.R., Allcock A.L. 2023. Genome skimming elucidates the evolutionary history of Octopoda. Mol. Phylogen. Evol. 182.

Talavera G., Castresana J. 2007. Improvement of phylogenies after removing divergent and ambiguously aligned blocks from protein sequence alignments. Syst. Biol. 56:564–577.

Team R.C. 2019. R: A language and environment for statistical computing. Vienna, Austria, R Foundation for Statistical Computing.

Tin M.M.-Y., Economo E.P., Mikheyev A.S. 2014. Sequencing degraded DNA from non-destructively sampled museum specimens for RAD-tagging and low-coverage shotgun phylogenetics. PLoS ONE 9:e96793.

Van Dam M.H., Lam A.W., Sagata K., Gewa B., Laufa R., Balke M., Faircloth B.C., Riedel A. 2017. Ultraconserved elements (UCEs) resolve the phylogeny of Australasian smurf-weevils. PLoS ONE 12:e0188044.

Wickham H. 2016. ggplot2: Elegant Graphics for Data Analysis. New York, Springer-Verlag.

Wong T.K.F., Ly-Trong N., Ren H., Baños H., Roger A.J., Susko E., Bielow C., Maio N.D., Goldman N., Hahn M.W., Huttley G., Lanfear R., Minh B.Q. 2025. IQ-TREE 3: Phylogenomic inference software using complex evolutionary models. EcoEvoRxiv.

Wood H.M., Gonzalez V.L., Lloyd M., Coddington J., Scharff N. 2018. Next-generation museum genomics: Phylogenetic relationships among palpimanoid spiders using sequence capture techniques (Araneae: Palpimanoidea). Mol Phylogenet Evol 127:907–918.

Young A.D., Gillung J.P. 2020. Phylogenomics — principles, opportunities and pitfalls of big-data phylogenetics. Syst. Entomol. 45:225–247.

Zeng C.X., Hollingsworth P.M., Yang J., He Z.S., Zhang Z.R., Li D.Z., Yang J.B. 2018. Genome skimming herbarium specimens for DNA barcoding and phylogenomics. Plant Methods 14:43.

Zhang F., Ding Y., Zhu C.D., Zhou X., Orr M.C., Scheu S., Luan Y.X., Matschiner M. 2019a. Phylogenomics from low-coverage whole-genome sequencing. Methods in Ecology and Evolution 10:507–517.

Zhang Y.M., Lucky A., Williams J.L. 2019b. Understanding UCEs: a comprehensive primer on using ultraconserved elements for arthropod phylogenomics. Insect Systematics and Diversity 3:1–12.

