## Supplemental Figures for "Low-Coverage Genome Sequencing Outperforms Target Enrichment Phylogenomics"

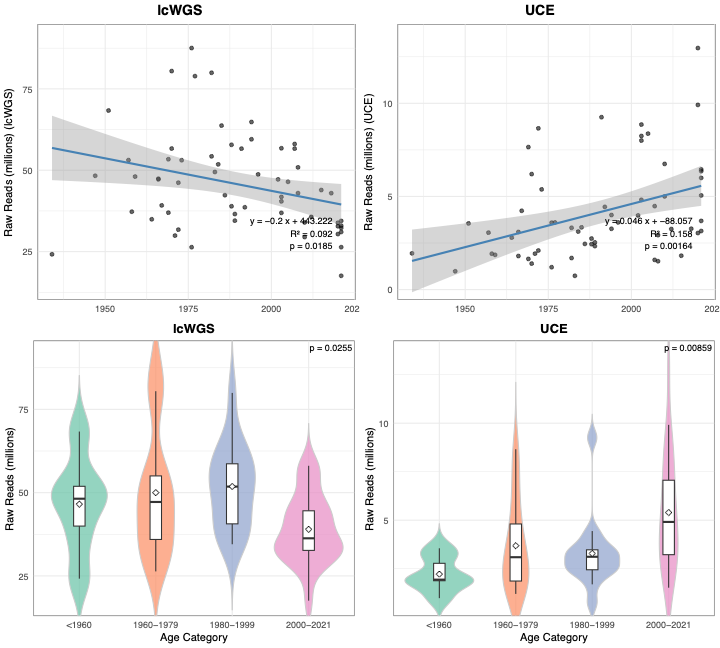

**Figure S1.** Relationship between collection year and raw paired-end read count for the lcWGS and UCE datasets. Upper graphs display scatter plots of read counts versus collection year with all samples data combined. Lower graphs are violin plots of read count distributions with samples grouped by age category.

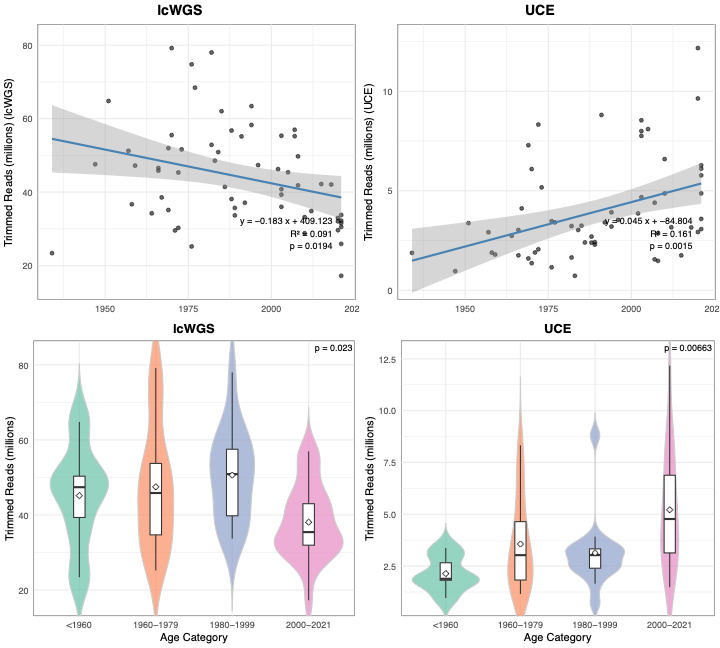

**Figure S2.** Relationship between collection year and trimmed paired-end read count for the lcWGS and UCE datasets. Upper graphs display scatter plots of read counts versus collection year with all samples data combined. Lower graphs are violin plots of read count distributions with samples grouped by age category.

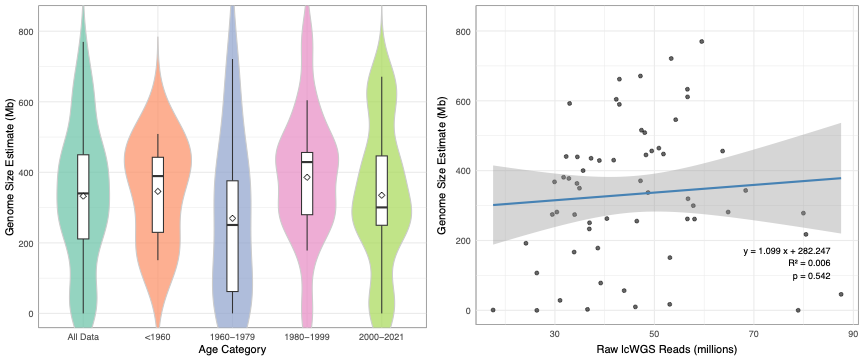

**Figure S3**. GenomeScope2 genome size estimates for all samples. The left plot shows size distributions for all data and by age category. The right plot shows genome size versus the raw read count and indicates no significant relationship.

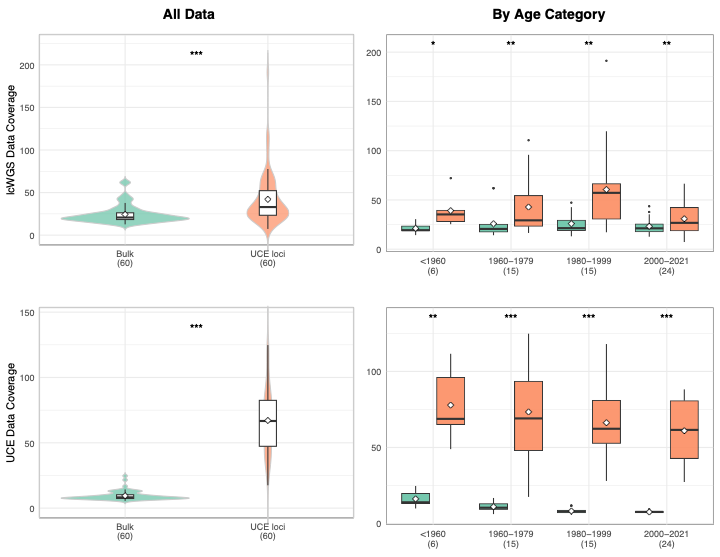

**Figure S4.** Sequencing coverage for lcWGS and UCE datasets comparing all assembled contigs (bulk) and uce contigs only. Top row shows results for lcWGS data comparing all data and results within age categories. Botton row shows results for the UCE-enriched dataset. Asterisks indicate level of significance between bulk and UCE contig coverage.

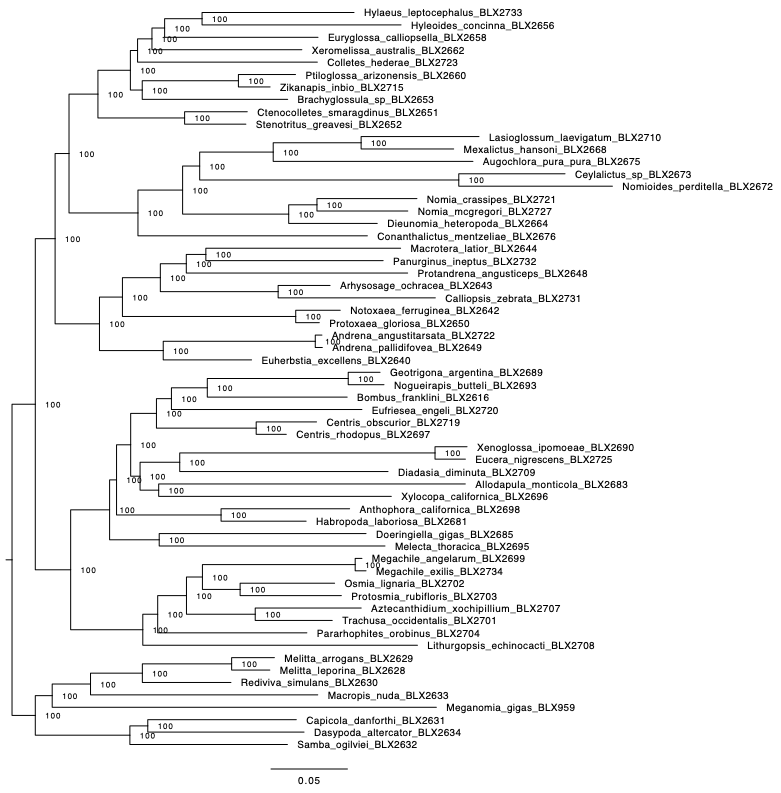

**Figure S5.** Concatenated Hymenoptera 2.5Kv2 UCE phylogeny based on the lcWGS dataset. Tree topology inferred using IQ-TREE v2 (GTR+F+G4). All nodes received 100% ultrafast bootstrap (UFB) support.

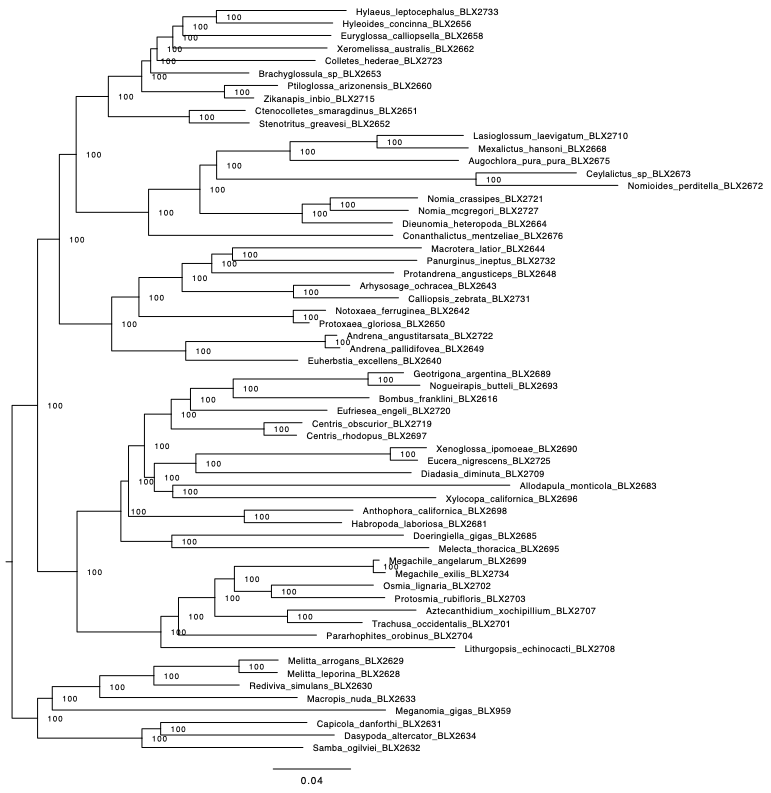

**Figure S6.** Concatenated Hymenoptera 2.5Kv2 UCE phylogeny based on the UCE-enriched dataset. Tree topology inferred using IQ-TREE v2 (GTR+F+G4). All nodes received 100% ultrafast bootstrap (UFB) support.

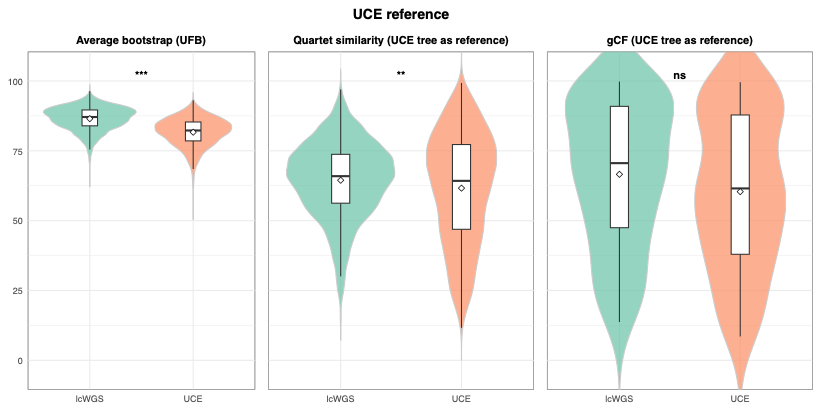

**Figure S7.** Comparison of gene tree performance metrics between lcWGS and UCE-enriched datasets. Violin and box plots show (left to right): mean ultrafast bootstrap (UFB) support across gene trees, quartet similarity to the UCE reference topology (gene tree “accuracy”), and gene concordance factors (gCF) across reference tree nodes (n = 57 nodes). Statistical significance was assessed using Wilcoxon signed-rank tests with multiple-testing correction where applicable. Asterisks denote significance levels (*** p < 0.0001; ns = not significant).

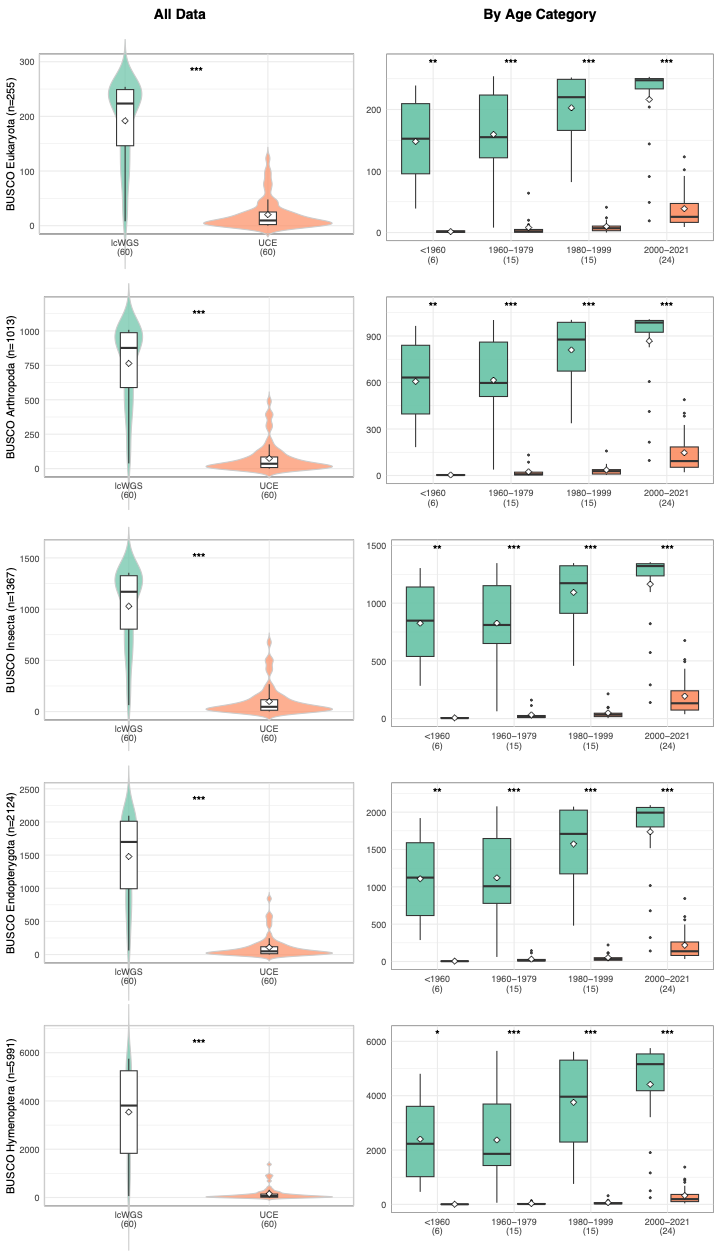

**Figure S8.** Recovery of BUSCO gene sets from lcWGS and UCE-enriched datasets. Left column shows all samples combined; right column shows results by age category. BUSCO gene sets from top to bottom are Eukaryota, Arthropoda, Insecta, Endopterygota, and Hymenoptera. Violin and box plots summarize distributions. Paired statistical comparisons between lcWGS and UCE datasets were performed per sample (Wilcoxon signed-rank or paired t-test as appropriate; BH-adjusted p-values). Asterisks indicate significance levels. All comparisons are paired by specimen.

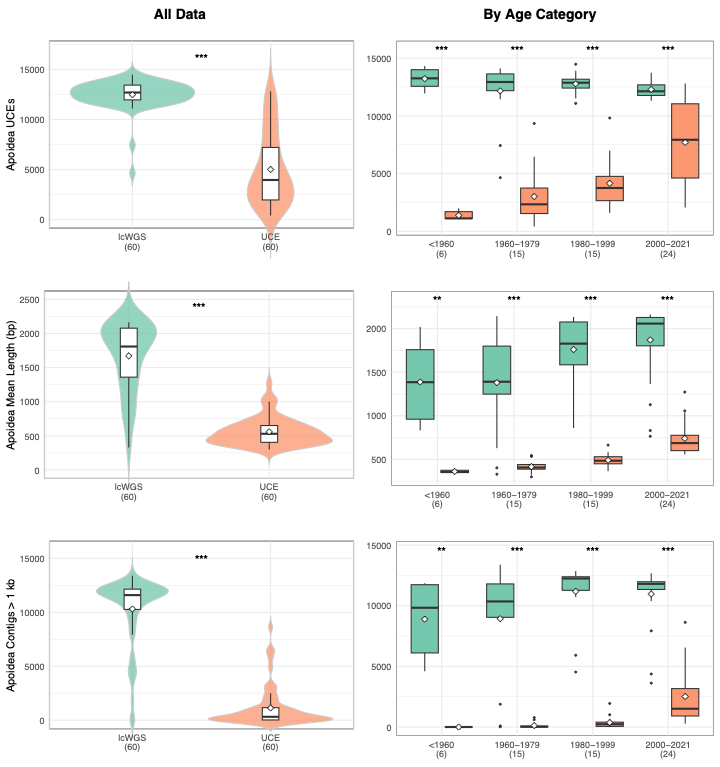

**Figure S9.** Comparison of Apoidea 20Kv1P UCE recovery metrics between lcWGS and UCE-enriched datasets. Left column shows all samples combined; right column shows results separated by age category. From top to bottom: (1) number of extracted UCE loci, (2) mean UCE locus length (bp), and (3) number of UCE contigs >1 kb. Violin plots show distribution density, boxplots show medians and interquartile ranges. Asterisks denote significance of paired comparisons between lcWGS and UCE datasets (Wilcoxon signed-rank or paired t-test as appropriate; Benjamini–Hochberg adjusted p-values): * p < 0.05, ** p < 0.01, *** p < 0.001. Sample sizes (n) are indicated on the x-axis. All comparisons are paired by sample.

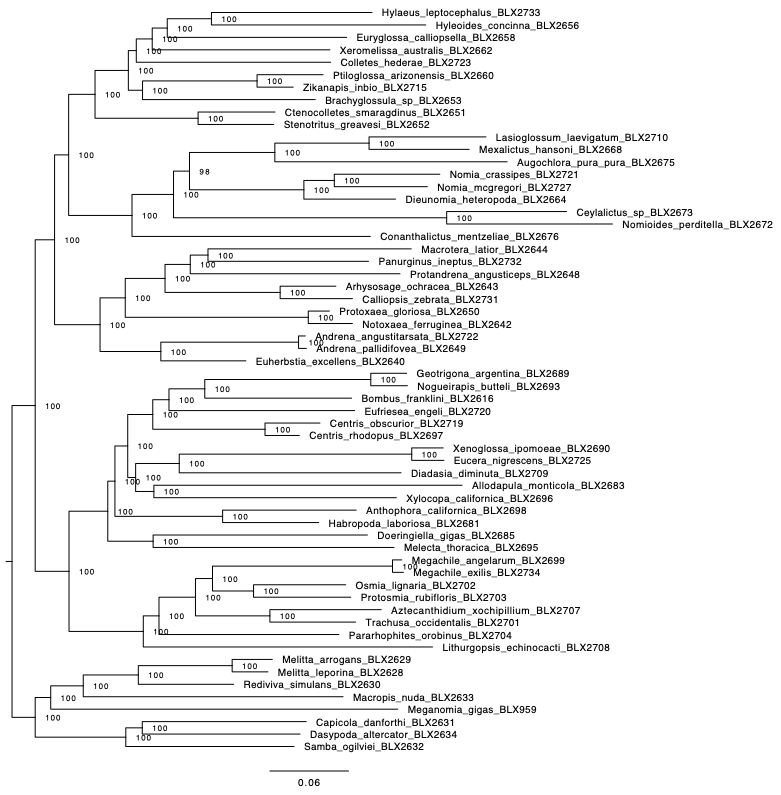
**Figure S10.** Concatenated phylogeny based on Apoidea 20Kv1P UCE loci extracted from the lcWGS dataset. Tree topology inferred using IQ-TREE v2 (GTR+F+G4). All nodes received 100% ultrafast bootstrap (UFB) support.
