## Supplemental Methods for "Low-Coverage Genome Sequencing Outperforms Target Enrichment Phylogenomics"

**Supplemental Methods: Computational Commands**

**Overview**

This section provides more detailed computational workflows and representative command-line examples used for processing low-coverage whole genome sequencing (lcWGS) and ultraconserved element (UCE) datasets. All analyses were conducted using standardized pipelines.

**Read Processing**

This section explains how raw reads were demultiplexed, trimmed, quality checked, and assembled.

- Raw reads were demultiplexed using BBTools v39.00 (lcWGS only).

demuxbyname2.sh prefixmode=f in=<R1.fastq.gz> in2=<R2.fastq.gz> out=<sample>_R1.fastq.gz out2=<sample>_R2.fastq.gz outu=Undetermined_R1.fastq.gz outu2=Undetermined_R2.fastq.gz names=my_barcodes.txt stats=read-counts.txt

- Raw reads were quality checked using FastQC v0.12.1.

fastqc -t <cores> --noextract --nogroup -o fastqc *.fastq.gz

- Raw reads were trimmed with Illumiprocessor v2.10.

illumiprocessor --input ./raw-fastq --output ./uce-clean --config ./illumiprocessor.conf --cores 48 --r1-pattern _R1.fastq.gz --r2-pattern _R2.fastq.gz

- Trimmed reads were quality checked using FastQC v0.12.1.

fastqc -t <cores> --noextract --nogroup -o fastqc *.fastq.gz

- Trimmed reads were assembled with SPAdes v3.14.1.

phyluce_assembly_assemblo_spades --output ./spades --cores <cores> --subfolder split-adapter-quality-trimmed --config assemblo.conf --memory <memory>

- Sequencing coverage was estimated using the Phyluce mapping workflow.

phyluce_workflow --config <config file> --output <output folder> --workflow mapping --cores <cores>

- For two samples, reads were normalized to 40x coverage prior to assembly using bbnorm.sh.

bbnorm.sh in1=R1.fastq.gz in2=R2.fastq.gz out1=R1.norm.fastq.gz out2=R2.norm.fastq.gz target=40 min=2 threads=<cores>

**Identification and Extraction of UCE Loci (lcWGS Only)**

This section explains how UCE loci were identified and extracted from the lcWGS assemblies.

- The Hymenopter 2.5Kv2P probes were aligned to the SPAdes assemblies using Phyluce.

phyluce_probe_run_multiple_lastzs_sqlite --db bees-phyluce-spades-60t.sqlite --output bees-phyluce-spades-60t-lastz --scaffoldlist <scaffold list> --genome-base-path ./genomes/ --probefile hymenoptera-v2-PRINCIPAL-bait-set.fasta --coverage 80 --identity 80 --cores <cores>

- 1kb of flanking DNA was sliced out from each identified UCE locus using Phyluce.

phyluce_probe_slice_sequence_from_genomes --lastz bees-phyluce-spades-60t-lastz --conf genomes.conf --flank 1000 --name-pattern "hymenoptera-v2-PRINCIPAL-bait-set.fasta_v_{}.lastz.clean" --output bees-phyluce-spades-60t-fasta-flank1000

**Processing UCE Contigs for Phylogenetics**

This section explains how UCE contigs were identified and processed for phylogenetics.

- The Hymenopbera 2.5Kv2P probe file was used to extract UCE contigs and remove potential paralogs using Phyluce.

phyluce_assembly_match_contigs_to_probes --contigs ./spades/contigs --probes ./ hymenoptera-v2-PRINCIPAL-bait-set.fasta --output ./lastz --min-coverage 80 --min-identity 80

- The number of UCE hits was summarized using Phyluce.

phyluce_assembly_get_match_counts --locus-db ./lastz/probe.matches.sqlite --taxon-list-config ./datasets.conf --taxon-group samples --output ./match.out --incomplete-matrix

- A monolithic FASTA file was generated based on the match results using Phyluce.

phyluce_assembly_get_fastas_from_match_counts --contigs ./spades/contigs --locus-db ./lastz/probe.matches.sqlite --match-count-output ./match.out --incomplete-matrix ./incomplete.out --output ./fastas.fasta

- The individual UCE loci were separated and aligned with MAFFT using Phyluce.

phyluce_align_seqcap_align --input ./fastas.fasta --output ./alignments/align-fasta --taxa 60 --aligner mafft --output-format fasta --incomplete-matrix --no-trim --cores <cores>

- Alignments were trimmed with Gblocks using Phyluce.

phyluce_align_get_gblocks_trimmed_alignments_from_untrimmed --alignments ./alignments/align-fasta/ --output ./alignments/align-nexus-gblocks --input-format fasta --output-format nexus --b1 0.5 --b2 0.5 --b3 12 --b4 7 --cores <cores>

- Locus names were removed from FASTA headers using Phyluce.

phyluce_align_remove_locus_name_from_files --alignments ./alignments/align-nexus-gblocks/ --output ./alignments/align-nexus-gblocks-names/ --cores <cores>

- Alignments were filtered for 75% taxon occupancy using Phyluce.

phyluce_align_get_only_loci_with_min_taxa --alignments ./alignments/align-nexus-gblocks-names/ --taxa 60 --output ./alignments/align-nexus-gblocks-names-75 --percent 0.75 --cores <cores>

- Alignments were concatenated for phylogenetic analysis using Phyluce.

phyluce_align_concatenate_alignments --alignments ./alignments/align-nexus-gblocks-names-75/ --output ./phylip/bees-60t-75p --input-format nexus –phylip

**UCE Statistics Generation**

This section explains how stats were generated for the UCE loci from each dataset.

- The monolithic FASTA file was exploded by taxon using Phyluce.

phyluce_assembly_explode_get_fastas_file --input ./fastas.fasta --output ./uce-contigs-exploded --by-taxon

- Statistics for each locus were generated using Phyluce.

phyluce_assembly_get_fasta_lengths --input sample.fasta --csv

**Concatenated IQ-TREE Analysis**

- The phylogenetic analyses for all concatenated datasets were performed with IQ-TREE v2.3.5.

iqtree-2.3.5 -s alignment.phylip -m GTR+G -B 1000 -T AUTO

**Genetree IQ-TREE Analysis**

- The phylogenetic analyses for all genetrees were performed with IQ-TREE v2.3.5.

iqtree-2.3.5 -s alignment.nexus -m MFP -AICc -B 1000 -T 1

**Gene Concordance Factor (gCF) IQ-TREE Analysis**

iqtree3 -t concat.treefile --gcf loci.treefile --prefix concord

**Apoidea Probe Set Design**

This workflow follows a modified version of Phyluce tutorial IV (<https://phyluce.readthedocs.io/en/latest/tutorials/tutorial-4.html>). Important modifications include the exact genomes selected, the probe alignment settings, and the number of retained loci.

- A selection of genomes was downloaded from NCBI (see Table S4).

- Genomes were converted to 2bit format.

faToTwoBit sample.fasta sample.2bit

- Reads were simulated using ART v2.5.8.

art_illumina --paired --in ~/path/to/input/genome.fasta

--out prefix-of-output-file --len 100 --fcov 2 --mflen 200 --sdev 150

-ir 0.0 -ir2 0.0 -dr 0.0 -dr2 0.0 -qs 100 -qs2 100 -na

- The read files were merged and compressed.

touch sample-pe100-reads.fq

cat sample-pe100-reads1.fq > sample-pe100-reads.fq

cat sample-pe100-reads2.fq >> sample-pe100-reads.fq

gzip sample-pe100-reads.fq

- The base genome was prepped with stampy v1.0.31 (Lunter and Goodson 2011).

stampy.py --species="Macropis_europaea" --assembly="Macropis_europaea_GCA916610135" -G Macropis_europaea_GCA916610135 Macropis_europaea_GCA916610135.fasta

stampy.py -g Macropis_europaea_GCA916610135 -H Macropis_europaea_GCA916610135

- Reads were aligned to the base genome using stampy.

python2.7 stampy.py --maxbasequal 93 -g <base genome> -h <base genome> --substitutionrate=0.05 -t <cores> --insertsize=400 -M <reads> | samtools view -Sb - > sample-to-base.bam

- Unmapped reads were removed with samtools.

samtools view -h -F 4 -b <sample>-to-base.bam > <sample>-to-base-MAPPING.bam

- BAM files were converted to BED format.

bedtools bamtobed -i <sample>.bam -bed12 > `basename <sample>`.bed

- Converted BEDs were sorted.

bedtools sort -i <sample>.bed > <sample>.sort.bed

- Overlapping or nearly overlapping intervals were merged.

bedtools merge -i <sample>.sort.bed > <sample>.merge.bed

- Repetitive intervals were removed.

phyluce_probe_strip_masked_loci_from_set --bed <sample>.sort.merge.bed --twobit Macropis_europaea_GCA916610135.2bit --output <sample>.strip.bed --filter-mask 0.25 --min-length 80

- A config file to identify loci shared across taxa was created.

- Shared alignment intervals were identified.

phyluce_probe_get_multi_merge_table --conf bed-files.conf --base-taxon Macropis_europaea_GCA916610135 --output anthophila-to-Macropis_europaea_GCA916610135.sqlite

- The resulting table was queried to see results.

phyluce_probe_query_multi_merge_table --db anthophila-to-Macropis_europaea_GCA916610135.sqlite --base-taxon Macropis_europaea_GCA916610135

- UCE loci were extracted.

phyluce_probe_query_multi_merge_table --db anthophila-to-Macropis_europaea_GCA916610135.sqlite --base-taxon Macropis_europaea_GCA916610135 --output Macropis_europaea_GCA916610135+12.bed --specific-counts 12

- The FASTA sequence was extracted from the base genome.

phyluce_probe_get_genome_sequences_from_bed --bed Macropis_europaea_GCA916610135+12.bed –twobit Macropis_europaea_GCA916610135.2bit --buffer-to 160 --output Macropis_europaea_GCA916610135+12.fasta

- A temporary bait set was designed from the base taxon.

phyluce_probe_get_tiled_probes --input Macropis_europaea_GCA916610135+12.fasta --probe-prefix "uce-" --design apoidea-v1 --designer branstetter --tiling-density 3 --two-probes --overlap middle --masking 0.25 --remove-gc --output Macropis_europaea_GCA916610135+12.temp.probes

- Duplicate baits were removed.

phyluce_probe_easy_lastz --target Macropis_europaea_GCA916610135+12.temp.probes --query Macropis_europaea_GCA916610135+12.temp.probes --identity 50 --coverage 50 --output Macropis_europaea_GCA916610135+12.temp.probes-TO-SELF-PROBES.lastz

phyluce_probe_remove_duplicate_hits_from_probes_using_lastz --fasta Macropis_europaea_GCA916610135+12.temp.probes --lastz Macropis_europaea_GCA916610135+12.temp.probes-TO-SELF-PROBES.lastz --probe-prefix=uce-

- Baits were aligned to each genome.

phyluce_probe_run_multiple_lastzs_sqlite --probefile ../bed/Macropis_europaea_GCA916610135+12.temp-DUPE-SCREENED.probes --scaffoldlist <scaffold list> --genome-base-path ../genomes --identity 50 --cores 48 --db Macropis_europaea_GCA916610135+12.sqlite --output apoidea-genome-lastz

- Sequence around conserved loci was extracted from exemplar genomes.

phyluce_probe_slice_sequence_from_genomes --conf apoidea-genome.conf --lastz apoidea-genome-lastz --probes 180 --name-pattern "Macropis_europaea_GCA916610135+12.temp-DUPE-SCREENED.probes_v_{}.lastz.clean" --output apoidea-genome-fasta

- Loci that were identified consistently were put in a table.

phyluce_probe_get_multi_fasta_table --fastas ../apoidea-genome-fasta --output multifastas.sqlite --base-taxon Macropis_europaea_GCA916610135

phyluce_probe_query_multi_fasta_table --db multifastas.sqlite --base-taxon Macropis_europaea_GCA916610135

phyluce_probe_query_multi_fasta_table --db multifastas.sqlite --base-taxon Macropis_europaea_GCA916610135 --output Macropis_europaea_GCA916610135+12-back-to-12.conf --specific-counts 12

- The final bait set was created.

phyluce_probe_get_tiled_probe_from_multiple_inputs --fastas apoidea-genome-fasta --multi-fasta-output Macropis_europaea_GCA916610135+12-back-to-12.conf --probe-prefix "uce-" --designer branstetter --design apoidea-v1 --tiling-density 3 --overlap middle --masking 0.25 --remove-gc --two-probes --output apoidea-v1-master-probe-list.fasta

- Duplicate baits were removed.

phyluce_probe_easy_lastz --target apoidea-v1-master-probe-list.fasta --query apoidea-v1-master-probe-list.fasta --identity 50 --coverage 50 --output apoidea-v1-master-probe-list-TO-SELF-PROBES.lastz

phyluce_probe_remove_duplicate_hits_from_probes_using_lastz --fasta apoidea-v1-master-probe-list.fasta --lastz apoidea-v1-master-probe-list-TO-SELF-PROBES.lastz --probe-prefix=uce-

**MitoFinder Mitogenome Recovery**

- Mitochondrial genes were identified in SPAdes assemblies using MitoFinder v1.4.1.

mitofinder -j <sample name> -a <assembly> -r hymenoptera_mitogenomes_ncbi.gb -o 5 -p 8 -m 24 -e 0.001 -n 15 --min-contig-size 500 --blast-size 20 –ignore

**BUSCO Analysis**

- BUSCO genes were identified in SPAdes assemblies using BUSCO v5.

busco -i <sample>.contigs.fasta -o <sample>_hymenoptera -l busco_db/hymenoptera_odb10 -m geno -c 48 --offline
